# Marrow-mimicry MEW scaffolds recapitulate erythroblastic islands for high-density cRBC biomanufacturing

**DOI:** 10.64898/2026.10.23.761469

**Authors:** Gloria Astrid Nausa Galeano, Susana Alexandra Costa Maia, Naomi C. Paxton, Kavita Bisht, Yoel Garcia Marin, Ryan McKinnon, Chloe M. de Nys, Keyu Tao, Lorenzo Moroni, Justin J. Cooper-White, Maria A. Woodruff, Rose Ann Franco, Ashley R. Murphy, Rebecca E. Griffiths, Mark C. Allenby

## Abstract

Biomanufactured cell therapies are delivering a step-change in modern medicine, but their high costs have prevented widespread adoption. Cultured red blood cells (cRBCs) are one type of cell therapy in clinical trials for off-the-shelf blood transfusion but remain limited by the exorbitant costs of manufacturing trillions of RBCs required for a blood transfusion. These costs could be mitigated if culture platforms could grow cells at higher density, like those found in the bone marrow, requiring less media and lower concentrations of expensive growth factors per cell.

We engineered melt electrowritten (MEW) polycaprolactone (PCL) scaffolds in geometries which imitate marrow trabeculae structures and were then plasma-treated to enable peripheral blood mononuclear cells (PB MNC) attachment. PB MNC monocytes first adhered to the scaffold and rapidly differentiated into CD68^+^ macrophage-like cells. These macrophage-loaded scaffolds adhered CD71^+^ erythroblasts, which then grew from 10 to 25% between days 4 and 8 and produced CD235a^+^ erythroid cells by day 20, under erythrogenic culture conditions. MEW PCL scaffolds in trabecular-like geometries had greater surface area, adhered more macrophages, and generated more erythroblasts than traditional MEW crosshatch geometries. The MEW scaffolds continuously released less-adherent cells to the supernatant, providing a 74% increase in supernatant cell numbers versus liquid suspension controls with identical cell seeding numbers and media volumes. High-cell-density scaffolds exhibited increased metabolism and endogenous production of self-supportive cytokines, which may offer cost-saving opportunities through future media optimisation.

This study reports marrow trabeculae-mimetic MEW PCL designs capable of selectively adhering PB MN cells to develop macrophage-coated scaffolds, which provided physical and paracrine support for cRBC production. This high-cell-density culture platform may be useful for future cell therapy biomanufacturing protocols.

## 1. Introduction

Ex vivo erythropoiesis has emerged as a promising strategy to generate cultured red blood cells (cRBCs) for transfusion, demonstrated by early clinical proof-of-concept studies and ongoing trials comparing manufactured cells to donor-derived products [1–4]. Despite this progress, translation remains constrained by biological and technological limitations, particularly the high cost and inefficiency of current platforms.

Current clinical trials of cRBCs require up to US$50,000 worth of culture media to produce a microdose of cRBCs equivalent to 5%-10% of a full blood bag [5,6]. This high cost is a result of the large quantity of media and concentrated biomolecular factors required to manufacture cell and gene therapies, including cRBCs, in low-cell-density liquid suspension bioreactors. Furthermore, these systems typically yield reticulocytes rather than fully mature erythrocytes, with overall expansion limited by the finite self-renewal capacity of erythroid progenitors and by cell loss during differentiation [7].

After decades of optimisation of liquid suspension erythropoiesis systems, it has become increasingly apparent that many of their limitations arise from the inability to reproduce the local microenvironment of haematopoietic organs [8–10], which in vivo orchestrate the rightly regulated balance between haematopoietic stem cell (HSPC) and progenitor quiescence, expansion, and lineage commitment. To overcome the limitations of suspension systems, there has been growing interest in developing three-dimensional (3D) culture platforms capable of recapitulating key features of the bone marrow (BM) microenvironment [11–15].

Biomaterial-based scaffolds offer the possibility to integrate structural, mechanical, and biochemical cues governing haematopoiesis, including extracellular matrix (ECM) interactions, mechanotransduction pathways, and spatial organisation of cellular populations [16–18]. Porosity, pore size, and mechanical properties can be tuned to facilitate nutrient transport and cell migration while mimicking key architectural features of trabecular bone [19–22].

Among the available fabrication techniques, melt electrowriting (MEW) has emerged as a promising approach for engineering highly controlled fibrous scaffolds. MEW enables the precise deposition of microscale polymer fibres (often 2-50 µm in diameter), bridging the gap between electrospinning and conventional extrusion-based printing [23]. This high-resolution control allows the design of reproducible architectures with pore sizes in the range of hundreds of micrometres, making MEW particularly suitable for generating 2.5D scaffolds that mimic trabecular structures found within haematopoietic BM microenvironments [22].

Trabecular BM architectures provide a large surface area for cell adhesion while maintaining open porosity, thereby supporting both cell attachment and mass transport, even in the absence of an engineered blood supply [24]. Importantly, thin MEW polycaprolactone (PCL) scaffolds offer an intermediate level of complexity between two-dimensional cultures and fully 3D constructs, enabling the investigation of early cell-matrix interactions without the diffusion limitations often associated with dense 3D systems. This makes MEW scaffolds especially relevant for studying and screening conditions supportive of erythropoiesis, where cell adhesion, viability, and spatial organisation are critical determinants of outcome.

Furthermore, the MEW PCL scaffolds used in this study did not require extracellular matrix coatings, animal-derived proteins, or feeder-layer support to promote cell attachment. The use of a fully synthetic and reproducible biomaterial platform is advantageous from a translational perspective, as it reduces biological variability and simplifies future implementation in Good Manufacturing Practice (GMP)-compliant manufacturing workflows [25,26].

In this context, the comparison between MEW-based scaffold cultures and conventional liquid suspension systems is essential to evaluate the added value of biomimetic architectures. While suspension cultures provide homogeneous conditions and remain the benchmark for erythroid expansion, they lack the physical and cellular cues of the BM microenvironment. In contrast, MEW scaffolds can support localised cell-cell interactions and may reduce the dependence on exogenous growth factors by promoting endogenous signalling within the cultured cell populations [8,27].

To further recreate some of the cellular complexity of the BM microenvironment, this study employs peripheral blood mononuclear cells (PB MNCs) as the starting population within the 2.5D MEW-based culture system. PB MNCs comprise a heterogeneous mixture of 45-70% T-cells, 10-30% monocytes, and 10-20% of other cells including B-cells, NK cells, and additional leukocytes, and roughly 1% haematopoietic stem and progenitor cells (HSPCs). Although purified CD34^+^ HSPCs from PB, BM, or umbilical cord blood are often used as the primary starting population for ex vivo erythroid cultures, PB supports the development of an autologous and more accessible source for cell therapy manufacturing versus BM and cord blood. Furthermore, several studies have shown that retaining the accessory populations within unselected or minimally manipulated PB MNC fractions can enhance erythroid expansion and maturation compared with CD34^+^-only cultures [7,28,29]. This advantage is attributed to the retention of CD34^−^ progenitors and supportive cell populations, as well as the preservation of endogenous cell-cell interactions. For instance, monocytes within the MNC fraction can differentiate into macrophages upon adhesion to scaffold fibres, providing a functional component of the haematopoietic niche [30–32].

The local cell-cell interaction between different cell populations is especially relevant in the context of erythropoiesis, where erythroblastic islands, specialised microenvironments composed of a central macrophage surrounded by differentiating erythroid cells, play a critical role [33–35]. These structures regulate erythroblast proliferation, enucleation, haemoglobinisation, and present and secrete adherence and growth factors required for erythroblast differentiation [36–39]. By leveraging the intrinsic cellular diversity of PB MNCs, the present model aims to partially recapitulate these niche-like interactions without the need for exogenous stromal co-culture systems, thereby reducing regulatory complexity.

Overall, the integration of diverse PB MNCs with MEW-fabricated trabecular scaffolds represents a promising approach for developing a 2.5D culture platform that captures key aspects of the BM microenvironment for autologous and accessible cell therapy manufacturing. By systematically evaluating the effects of scaffold architecture on cell adhesion, viability, differentiation, and culture dynamics, this approach seeks to investigate whether controlled MEW microarchitectures can support the formation of early erythropoiesis-supportive cellular niches in processed PB MNC populations.

## 2. Materials and Methods

### 2.1. Scaffold fabrication

22 x 22 mm MEW scaffolds were fabricated using a custom-built printer previously described [40,41]. Polycaprolactone (PCL, 45 kDa, 704105, Sigma-Aldrich, St. Louis, MO, USA) was loaded at 90°C and extruded through a 25G needle. A G-code for a 0/90° fibre orientation with a 300 µm pore size was generated and uploaded to the printer. Four translational speeds above and below critical speeds were used (800, 600, 400, and 200 mm min^−1^) and produced 3D scaffolds with four distinct crosshatched-to-trabecular architectures. Fibres were deposited onto a glass plate at a working distance of 3mm, with an applied voltage of 3.2 kV and an air pressure of 0.1 MPa.

### 2.2. Scaffold structural analysis

For structural analysis and cell culture, discs of 5 mm diameter were cut using a disposable biopsy punch. Each architecture was printed with a different number of layers to produce 2.5D structures with similar total volume, 4.77 mm^3^ ± 0.94. Scaffold thickness and individual fibre diameter (n=12) were measured from confocal microscopy images acquired at 10x magnification using a confocal laser scanning microscope (Leica TCS SP8, Leica Mycrosystems, Wetzlar, Germany) and analysed using Fiji distribution of ImageJ (version 1.54, National Institutes of Health, NIH, Bethesda, MD, USA) [42]. Micro-computed tomography (mico-CT) scans (n=3) were collected using a 225 kV directional microfocus X-ray tube at a spatial resolution of 7.26 ± 0.55 µm (FF35 CT, Comet Yxlon, Hamburg, Germany). Structural analyses were performed in ImageJ software using the BoneJ plugin (version 7.0.19, National Institutes of Health, Bethesda, MD, USA) [43] to quantify porosity, surface area and mean pore size for each scaffold (10 independent measurements per scaffold).

### 2.3. Scaffold preparation

A total of 24 scaffolds, 6 technical replicates of 4 different architectures, were used for cell culture experiments per donor, for a total of 3 PB donors. Scaffolds were biopsy-punched to a 5 mm diameter. The day before the experiment starting time, scaffolds were oxygen plasma treated for 90 seconds and then flipped over to be treated for an extra 90 seconds for surface etching to promote cell attachment (RF Power: 35 W, Pulse: 255, Oxygen: 5.0 sccm) (Tergeo plasma cleaner 200315T203, PIE Scientific LLC, CA, U.S.A.).

Immediately after oxygen plasma treatment, scaffolds were treated with 70% ethanol, UV light, and washed with sterile Dulbecco’s phosphate-buffered saline (DPBS; Cat. No. 14190144, Gibco, Thermo Fisher Scientific, Waltham, MA, USA) three times. After sanitisation, scaffolds were transferred to a fresh 96-well plate and conditioned in basal medium IMDM with glutaMAX (FG0465, Sigma-Aldrich) at 37°C with 5% CO_2_ overnight in an incubator.

Before cell seeding, the conditioned scaffolds were transferred to a 96-well plate whose wells were previously coated with polydimethylsiloxane (PDMS) discs to reduce PB MNCs attachment to the bottom of the well and promote MNCs interaction with the scaffolds. PDMS discs were produced by soft lithography using SYLGARD™ 184 Silicone Elastomer (Dow, Midland, MI, USA) mixed at a 10:1 ratio. PDMS layers were biopsy-punched to create discs of 6 mm in diameter that were sterilised by team sterilisation and were placed at the bottom of each well.

### 2.4. Erythropoiesis culture

Volume-reduced PB units were collected from three consenting healthy donors by the Australian Red Cross Lifeblood and then cultured in accordance with approved ethics and agreements (2022/HE001331). Unless otherwise specified, all chemical reagents and buffers in this section were purchased from Sigma-Aldrich (St. Louis, MO, USA). PB MNCs were then isolated by diluting PB units in Hank’s Balanced Salt Solution (HBSS; Cat. No. H6648) supplemented with 0.6% v/v citrate-dextrose solution (ACD; Cat. No. C3821) and separated by gradient centrifugation over Histopaque (Cat. No. H8889). After 30 minutes of centrifugation at 400 × g, the MNC fraction was collected and washed five times with HBSS+ACD. Each wash step was followed by centrifugation at 250 × g for 10 min. Erythroid cells were lysed with a custom-made lysis buffer containing 150 mM NH_4_Cl (Cat. No. A943), 1 mM K2 EDTA.2H_2_O (Cat. No. ED2P), 10 mM KHCO_3_ (Cat. No. 237205), and NaOH (Cat. No. S8045) to adjust the pH to 7.5. After 5 minutes, the lysis buffer solution was neutralised with HBSS+ACD and viable MNCs were calculated using Trypan Blue (Cat. No. 15-250-061).

Immediately after each isolation, 2 × 10^5^ PB MNCs were suspended in 100 µL of complete media (2 × 10^6^ MNCs mL^−1^) and seeded onto the scaffolds. Samples were kept in an incubator, and after four hours, an extra 100 µL of media was added (final seeding density of 1 x 10^7^ cells mL^−1^). Cells were cultured in complete media comprised of Iscove’s Modified Dulbecco’s Medium (IMDM; Cat. No. FG0465, Sigma-Aldrich) supplemented with 3% v/v Human AB serum (H4522), 200 µg mL^−1^ Holo-Transferrin (Cat. No. 2914-HT, R&D Systems, Minneapolis, MN, USA), 10 µg mL^−1^ Insulin (I-9278), 3 U mL^−1^ Heparin (H3149), 3 U mL^−1^ erythropoietin (EPO; Cat. No. 78007, STEMCELL Technologies, Vancouver, BC, Canada), 2 mg mL^−1^ human serum albumin (HSA; Cat. No. 186-0002-0, SeraCare Life Sciences, Milford, MA, USA), 10 ng mL^−1^ stem cell factor (SCF, 255-SC-050, R&D Systems), and 1 ng mL^−1^ interleukin-3 (IL-3, 203-IL-010, R&D Systems), based on a protocol described previously [44]. Additionally, a liquid suspension control was cultured in parallel with the scaffold cultures in similar 96-well plates, following the same cell seeding density and media replenishment volume and frequencies.

Early erythropoiesis cultures were maintained for 8 days in total, with media changes every 48 h. Two cell groups were identified: those attached to the scaffold and those growing in suspension in the proximity of the scaffold. During each media change, the culture media was removed, minimising scaffold disturbance. The medium was centrifuged, and the cell pellet was resuspended in fresh medium to be placed back on the scaffolds. Cell suspension number and viability were quantified during each media change using Trypan Blue exclusion. The culture was divided into two time points, day 4 and day 8. On the 4th day of culture, half the scaffolds were transferred to a new 48-well plate coated with PDMS at the bottom, increasing the volume to 0.8 mL to meet cell growth requirements.

The prototype complete erythropoiesis culture lasted for 20 days in total, with media changes every 48 h. The same media composition described above was provided from day 0-8. From day 10, stem cell growth factors were removed from the media (SCF, IL-3), and holo-transferrin was increased to 300 ng mL^−1^ to support erythroblast differentiation. The volume of the culture media per sample was increased across the culture from 200 µL at day 0 to 800 µL at day 4, 1 mL at day 8, and 1.6 mL at day 10.

### 2.5. Immunofluorescence microscopy

At days 4, 8, and 20, scaffolds were fixed using 3% formaldehyde (Cat. No. C006, ProSciTech, Kirwan, QLD, Australia), then stained with 0.1% w/v Sudan Black B (Cat. No. 199664, Sigma-Aldrich). Cells were permeabilised with 0.025% Triton X-100 (Cat. No. 93443, Sigma-Aldrich) and blocked with 10% goat serum (Cat. No. 16210064, Gibco).

Scaffolds were stained with DAPI (Image-iT™ ReadyProbes™, Cat. No. R37606, Invitrogen, Thermo Fisher Scientific), F-actin (Cat. No. R37110, Invitrogen), a fixable viability stain (Live-or-Dye™ 640/662, Cat. No. 32007, Biotium, Fremont, CA, USA), and primary antibodies, all from Invitrogen: CD45 (1 µg mL-1, Mouse IgG1k, Cat. No. 14-0459-82), CD71 (1 µg mL-1, Rabbit IgG, Cat. No. MA5-33038), CD235a (1 µg mL-1, Rat IgG2b, Cat. No. MA1-80131), CD68 (2.5 µg mL-1, Mouse IgG1k, Cat. No. 14-0688-82), HIF-1α (10 µg mL-1, Rabbit IgG, Cat. No. PA1-16601).

Scaffolds were analysed by confocal microscopy using an Evident Olympus FV3000 confocal laser scanning microscope (Evident Corporation, Tokio, Japan) employing a 20x objective and recording z-stacks composed of individual images taken every 2 µm. Five z-stacks were taken per sample in posterior, anterior, left, right, and centre regions of interest to get representative information on cell numbers and populations, as illustrated in figure S 2 and figure S 3.

### 2.6. Quantitation of F-actin coverage and attached-cell viability

For the cells attached to the scaffolds, viability was assessed after fixation at days 4 and 8, using Live-or-Dye viability stain. Images were acquired using a confocal laser scanning microscope (Leica TCS SP8, Leica Mycrosystems) and a 20x objective.

Both F-actin and viability stains were analysed in Fiji distribution of ImageJ (version 1.54, National Institutes of Health) based on the number of thresholded voxels of each corresponding channel. The percentage of dead cells was calculated as the ratio between the total number of thresholded voxels and the average number of voxels associated with a single nucleus.

### 2.7. Scaffold-attached cell quantitation

Scripts for nuclei and stain segmentation and tracking were designed employing TrackMate (version 7.11.1) and StarDist (0.4.0) plugins [45,46] within the Fiji distribution of ImageJ (version 1.54p, NIH) [42,47] and then nuclei quantitation and localisation across the 3D image using a Python script (version 3.11, Python Software Foundation, Wilmington, DE, USA). Briefly, nuclear segmentation was performed by applying the StarDist detector on the 3D z-stacks. For each z-slice, StarDist detected nuclei after the application of filters for minimum quality, maximum perimeter and minimum mean intensity across the object. Detected spots were represented as star-convex polygons defined by 32 boundary vertices. TrackMate subsequently linked corresponding detections across a minimum of four adjacent z-slices, with a maximum linking distance of 2 µm between two consecutive spots. TrackMate then generated 3D nuclear objects represented by unique nucleus IDs. Using a Python script, polygon coordinates corresponding to the slice-specific contours of each nucleus were exported and were used for downstream quality control, colocalisation and intensity analyses. Nuclear segmentation accuracy was calculated against manual scoring of images, the details of which are provided in Supplementary figure S 2.

### 2.8. Phenotypic stain co-localisation with segmented nuclei

The colocalisation strategy was based on the overlap between the nuclei polygons generated by TrackMate for every z-slice and the thresholded image of every independent marker. To define each threshold, every cell marker was associated with its specific isotype. Isotype images were analysed and colocalised with their respective nuclei, and the threshold was determined based on intensity distributions measured exclusively within cell-associated ROIs of isotype controls. ROIs were defined as the area overlapping the polygon nuclei and a ring of 5 pixels larger than the polygon. Then every independent channel was thresholded using the 99.9th percentile of intensity from its corresponding isotype and then colocalised with its nuclei polygons. Results are presented as the percentage of cells positive for each marker and volumetric and/or surface density. Details of this method and representative outputs are provided in Supplementary figure S 3.

### 2.9. Metabolic dynamics

Culture metabolic activity was evaluated by quantifying glucose consumption and lactate production in culture supernatants collected at days 4, and 8. Metabolite concentrations were measured by high-performance liquid chromatography (HPLC) using an LC-20 modular system (Shimadzu, Kyoto, Japan) equipped with a differential refractive index detector (Optilab dRI, Wyatt Technology, Santa Barbara, CA, USA).

Metabolite separation was performed using an ion-exclusion column and corresponding guard column (Rezex™ ROA-Organic Acid H+, Phenomenex, Torrance, CA, USA). Samples (50 µL) were analysed at a flow rate of 0.6 mL min^−1^ and a column temperature of 60°C employing 2.5 mM H_2_SO_4_ as the mobile phase. Chromatographic runs were conducted for 40 min per sample.

Glucose and lactate concentrations were determined by comparison with external calibration standards analysed under identical chromatographic conditions (Glucose, GA018, Chem-supply, Gillman, SA, Australia) (Lactic acid, L6661, Merck, Darmstadt, Germany). Representative chromatograms, calibration curves, and analytical parameters are provided in supplementary data. Metabolite consumption and production rates were calculated from concentration changes measured between consecutive media exchanges and normalised to the duration of the sampling interval.

### 2.10. Growth factors and inflammation markers

Key growth factors and inflammation markers were analysed in the supernatant recovered from cell culture at days 4 and 8. Samples were analysed by flow cytometry (CytoFLEX LX, Beckman Coulter, Brea, CA, USA) employing the LEGENDplex™ kit (BioLegend, Inc., San Diego, CA, USA) with pre-coated beads for EPO, SCF, Granulocyte Colony-Stimulating Factor (G-CSF), Interleukin-6 (IL-6), Interferon-γ (INF-γ), Tumour Necrosis Factor-α (TNF-α) following manufacturer protocols.

### 2.11. Statistical analysis

Statistical analysis was conducted using one-way analysis of variance (ANOVA) (for scaffold architecture characterisation) and two-way ANOVA for the rest of the analysis, with scaffold architecture and culture time as factors. The analysis was then followed by Tukey’s multiple comparisons test (α = 0.05) (version 11.0.0, GraphPad Prism, GraphPad Software, Boston, MA, USA).

## 3. Results

### 3.1. Trabecular MEW scaffold architectures can be fabricated with sub-critical translational printing speeds

MEW was employed to fabricate scaffolds using identical printing parameters except for translational speed. While all scaffolds were generated from the same base geometry, decreasing translational speed altered the final architecture through changes in fibre deposition behaviour. This produced a set of MEW PCL scaffold architectures with varying porosities, available surface areas, and mean pore sizes, which were used to investigate their ability to support the adhesion of PB MNC populations.

Representative confocal microscopy and micro-CT reconstructions revealed a transition from highly ordered square pores at 800 mm min^−1^ towards increasingly disordered fibre arrangements when translational speed decreased (figure 1 (A-B)). At speeds below the critical translation speed, the high voltage applied between the nozzle and collector caused the molten filament to oscillate, producing an anisotropic architecture in which the likelihood of adjacent fibres fusing increases. Furthermore, because air pressure was kept constant, reducing the translational speed led to a higher material output from the needle, thereby affecting all the parameters analysed.

**Figure 1.**
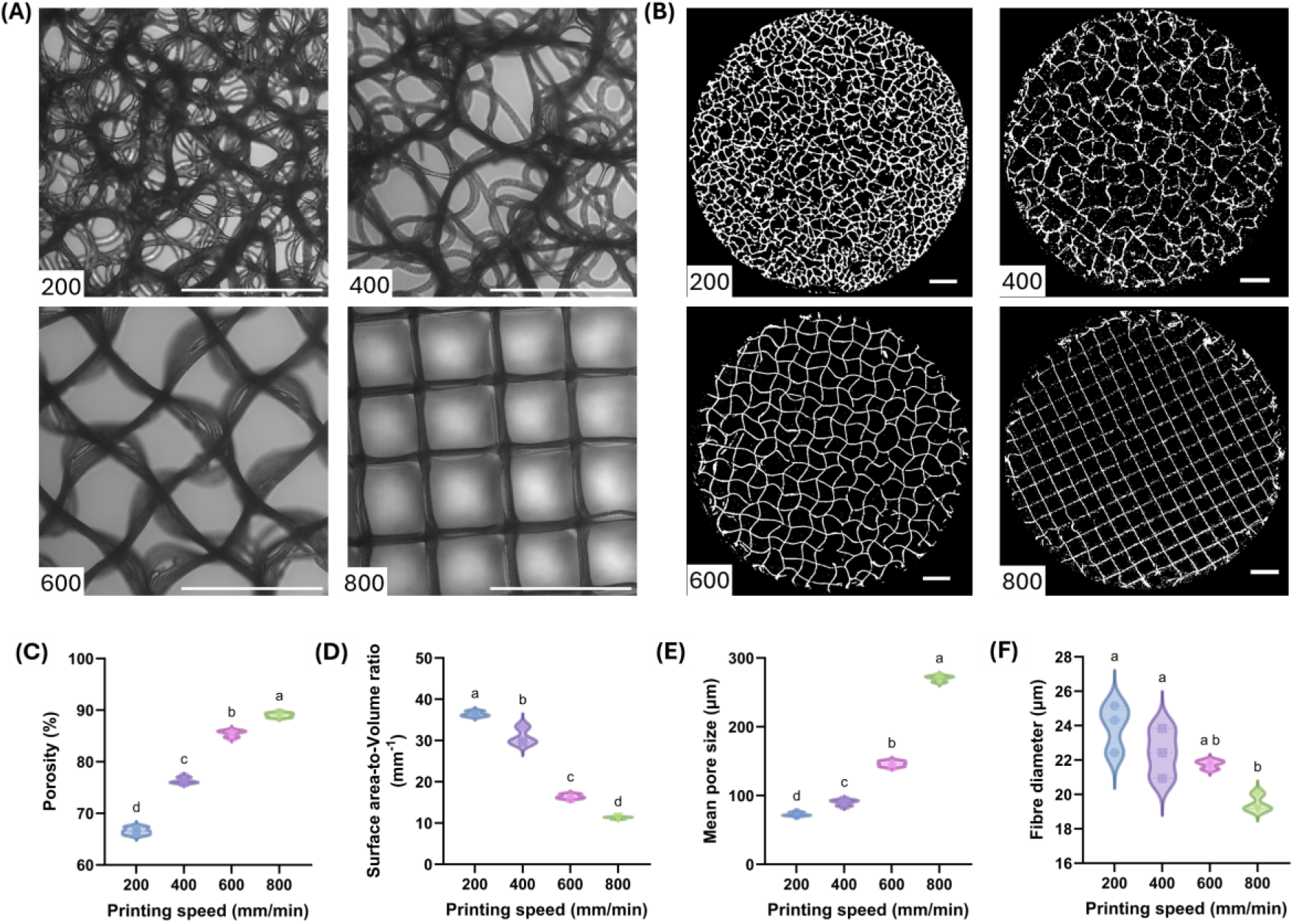
Structural characterisation of melt electrowritten (MEW) scaffolds. Scaffolds were fabricated at different translational speeds (200, 400, 600, and 800 mm min^−1^). (A) Representative confocal microscopy images showing scaffold morphology. (B) Representative micro-computed tomography (micro-CT) reconstructions used for structural analysis. Quantitative analysis of (C) porosity, (D) surface area-to-volume ratio, (E) mean pore size, and (F) fibre diameter. Data are presented as distributions. n=3. Groups sharing different letters are significantly different (p < 0.05). Scale bar: 500 µm.

Quantitative structural analysis confirmed that translational speed significantly influenced scaffold microarchitecture. Porosity and mean pore size increased with printing speed, while surface area-to-volume ratio and fibre diameter decreased. These observations agree with previous reports describing the transition around the critical translation speed in MEW, where fibre buckling and coiling become more pronounced below the collector matching velocity [48,49].

### 3.2. Scaffold architecture regulates attachment and spatial organisation of PB MNCs

A significant amount of PB MNCs readily attached to all scaffold architectures and remained viable throughout culture. Confocal imaging demonstrated substantial differences in cell organisation depending on scaffold morphology (figure 2 (A)). Cells cultured on low-speed architectures formed dense cellular aggregates associated with the highly entangled fibre networks, whereas cells cultured on the crosshatched 800 mm min^−1^ scaffolds showed preferential localisation along individual fibres.

**Figure 2.**
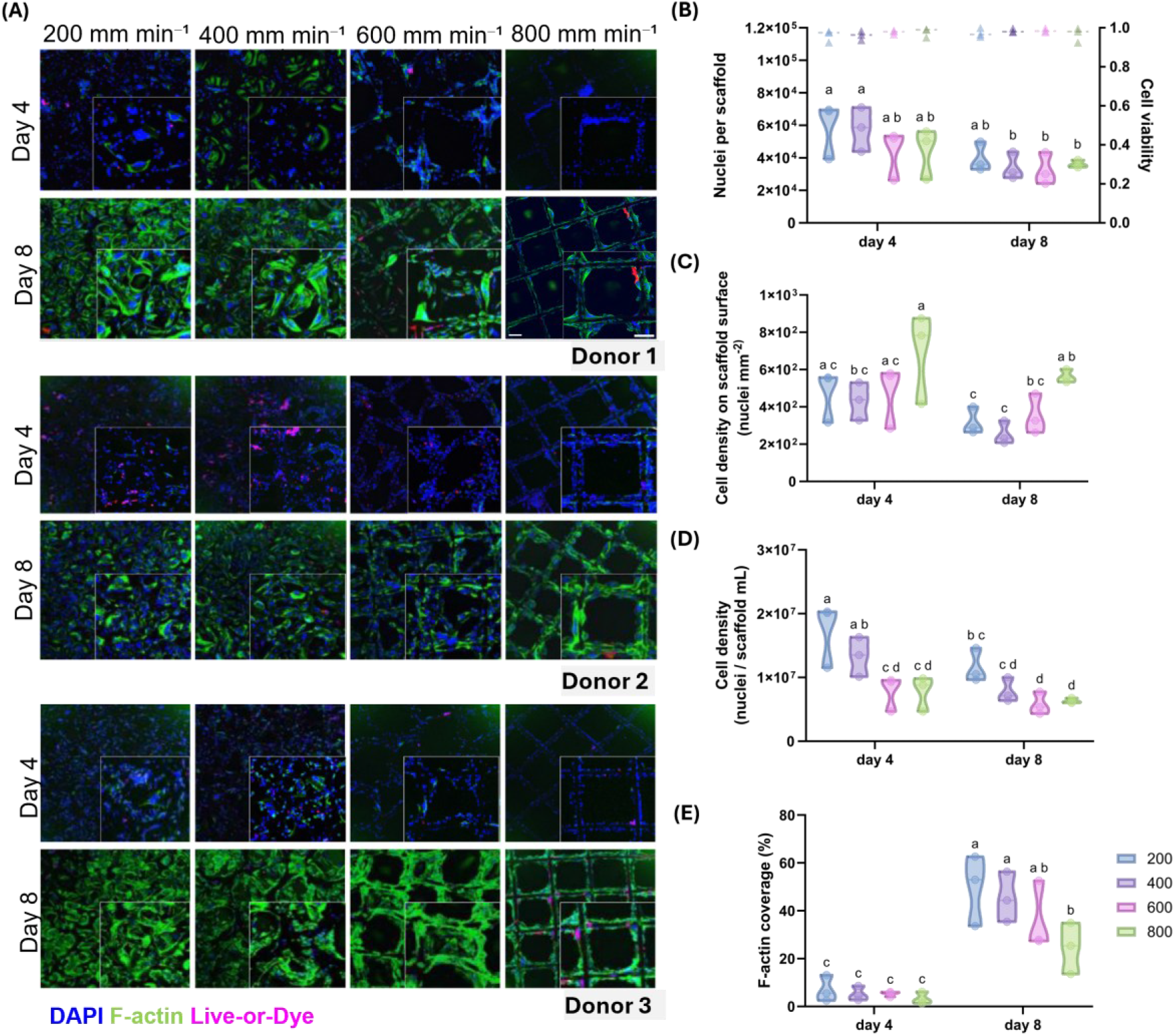
MEW scaffold architecture influences MNC adhesion, viability, and cytoskeletal organisation. (A) Representative maximum-intensity projections of confocal microscopy images acquired after 4 and 8 days of culture on melt electrowritten (MEW) scaffolds fabricated at translational speeds of 200, 400, 600, and 800 mm min^−1^. Cell nuclei, F-actin cytoskeleton, and viable cells were stained wih DAPI (blue), phalloidin (green), and Live-or-Dye viability stain (magenta), respectively. Scale bars: 100 µm. (B) Quantification of cell attachment and viability following fixation, fluorescent staining, confocal imaging, and image analysis. Violin plots show the distribution of nuclei counts per scaffold, and triangles indicate cell viability (%). (C) Cell density normalised to scaffold surface area (nuclei mm^−2^). (D) Cell density normalised to scaffold volume (nuclei mm^−3^). (E) Percentage of scaffold surface covered by F-actin. Data were obtained from three independent donors (n = 3). Groups sharing different letters are significantly different (p < 0.05).

Quantitative image analysis revealed significant architectural effects on the number of attached cells, cell density, and cytoskeletal coverage (figure 2 (B-E)). While viability remained high across all groups (97.4 ± 2.1%), scaffold architecture strongly influenced the extent of cell attachment and spreading. The largest pore architectures generally exhibited reduced cellular occupancy despite maintaining viability. These findings suggest that scaffold geometry primarily affects cell retention and organisation rather than survival itself.

Interestingly, a statistically significant decrease in the number of scaffold-associated nuclei was observed between days 4 and 8 despite viability remaining above 90% across all scaffold architectures (figure 2. B). This behaviour suggests that the scaffold-associated population is not highly proliferative, or that scaffold surfaces may have approached saturation, limiting the ability of cells to continue expanding.

Normalisation of cell number to scaffold surface area (figure 2 (C)) revealed that attachment efficiency did not directly correlate with the total available surface. Although the 200 mm min^−1^ architecture exhibited the largest surface area, this parameter alone was not sufficient to explain the distribution of attached cells across the different scaffold designs. These findings suggest that cell attachment is governed not only by the quantity of available surface but also by the spatial organisation of the pore architecture, as reported previously [19–21].

Cell density normalised to scaffold volume was highest in the 200 mm min^−1^ architecture at both culture time points, indicating that cells occupied the available volume more densely than in the more open scaffold designs (figure 2 (D)). The maximum cell density achieved within the scaffold volume was approximately 2 × 10^7^ cells mL^−1^. Such high cell densities are particularly relevant in the context of BM engineering because the native haematopoietic niche is characterised by densely packed cellular microenvironments that facilitate extensive cell-cell communication and local paracrine signalling [10,50].

Finally, F-actin coverage was quantified to investigate how cells physically interact with scaffold surfaces. A marked increase in F-actin coverage was observed between days 4 and 8 (figure 2 (E)). Because the total number of scaffold-associated nuclei decreased during the same period, the increase in F-actin signal suggests substantial cytoskeletal remodelling and cell spreading following scaffold attachment.

### 3.3. MEW fibres act as preferential adhesion sites for macrophage-like cells

Representative confocal images revealed CD68-positive macrophages significantly covered MEW scaffold fibres at days 4 and 8 (figure 3 (A)), representing approximately 70-90% of all attached cells (figure 3 (C)). At day 4, most CD68-positive cells displayed a rounded morphology and were distributed intermittently along fibres. By day 8, these cells exhibited extensive spreading and elongation, frequently extending along multiple fibre segments and spanning large portions of the scaffold architecture.

**Figure 3.**
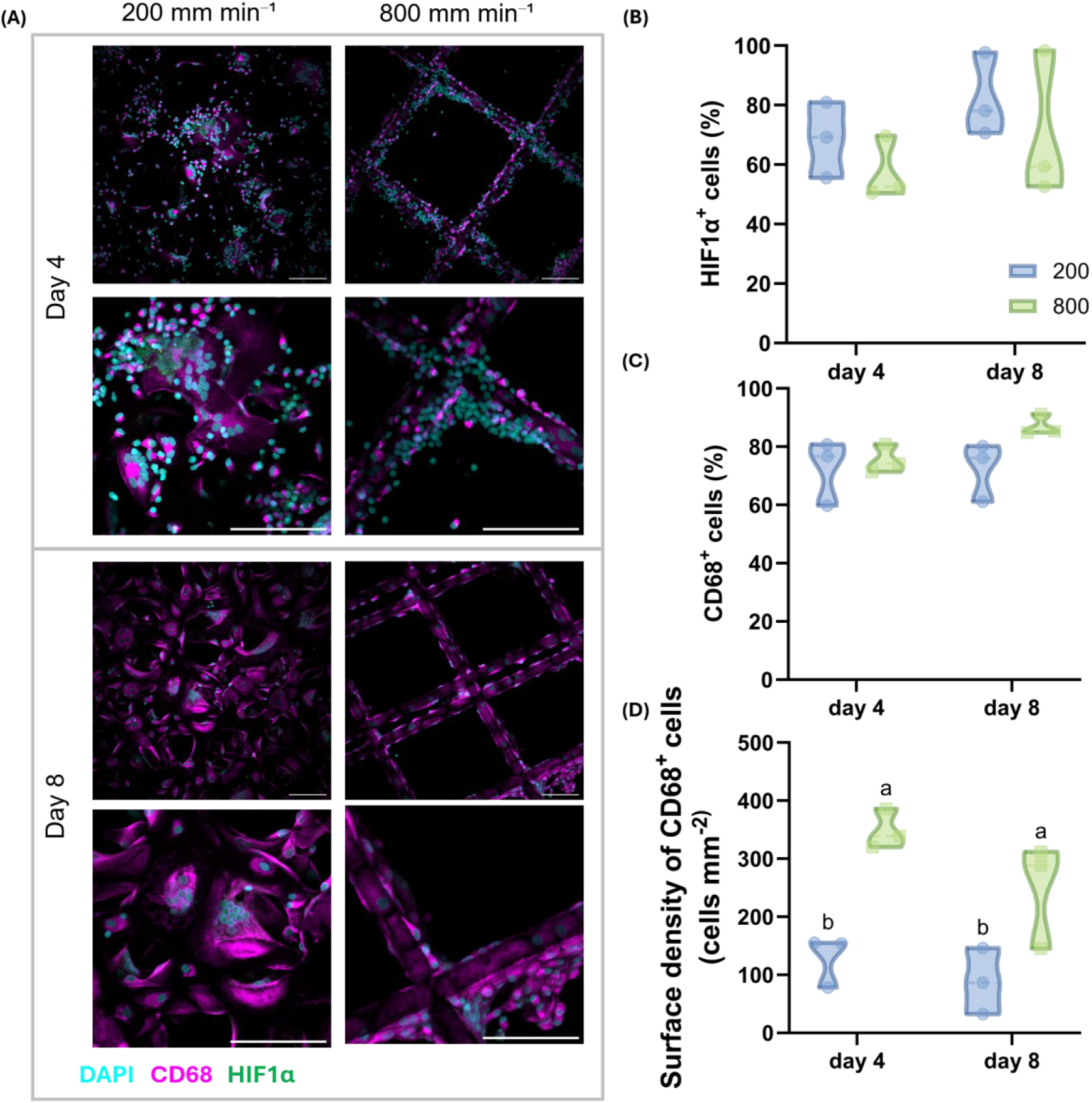
MEW scaffolds act as anchors for macrophage (CD68) adhesion. (A) Representative maximum-intensity projections of confocal microscopy images from Donor 1 acquired after 4 and 8 days of culture on melt electrowritten (MEW) scaffolds fabricated at translational speeds of 200 and 800 mm min^−1^. Cells were stained with DAPI (cyan), CD68 (magenta), and HIF1α (green). Scale bar = 100 µm. (B) Percentage of HIF1α-positive cells relative to the total number of nuclei. (C) Percentage of CD68-positive cells relative to the total number of nuclei. (D) Surface density of CD68-positive cells expressed as the number of CD68-positive cells per scaffold surface area (cells mm^−2^). Data were obtained from three independent donors (n = 3). Groups sharing different letters are significantly different (p < 0.05).

Surprisingly, neither culture duration nor scaffold architecture significantly altered the proportion of CD68-positive cells. This stability suggests that macrophage-like cells are not a transient population progressively replaced by erythroid progenitors but rather represent a persistent component of the scaffold-associated microenvironment. Additionally, considering that cultures were maintained under conditions optimised for erythroid proliferation rather than macrophage expansion, the sustained prevalence of CD68-positive cells likely reflects efficient retention and survival of these cells on scaffold fibres.

Despite displaying similar percentages of CD68-positive cells, macrophage density normalised to scaffold surface area was substantially greater in the 800 mm min^−1^ architecture (figure 3 (D)). This finding suggests that scaffold organisation influences the spatial distribution of macrophages more strongly than their relative abundance. Confocal imaging revealed that CD68-positive cells progressively elongated along fibre surfaces, particularly within the ordered architectures, indicating that macrophages actively utilised scaffold fibres as anchoring and spreading substrates. Consequently, highly ordered architectures appear particularly effective at promoting macrophage occupancy of the available scaffold surface.

Additional evidence of niche remodelling was obtained through phenotypic characterisation of HIF-1α expression. HIF-1α-positive cells were detected throughout the culture period, indicating activation of pathways commonly associated with hypoxic adaptation, metabolic remodelling, and inflammatory signalling (figure 3 (B)).

### 3.4. Macrophage-loaded trabecular architectures support greater erythroid commitment

Representative confocal images revealed the presence of abundant CD45-positive cells across all scaffold architectures and culture time points (figure 4 (A)). Two distinct populations could be identified based on their morphology and spatial localisation. First, large CD45-positive cells were frequently observed attached directly to scaffold fibres, with progressive spreading, elongation, and flattening against fibres between days 4 and 8, likely to be CD45-positive and CD68-positive macrophages. Second, small rounded CD45-positive cells were observed within the pore spaces and in close proximity to the fibres without any clear polarisation. The coexistence of these two populations suggests that scaffold cultures contain both adherent and suspension-associated haematopoietic cells, consistent with a dual-compartment culture system described in figure 6.

**Figure 4.**
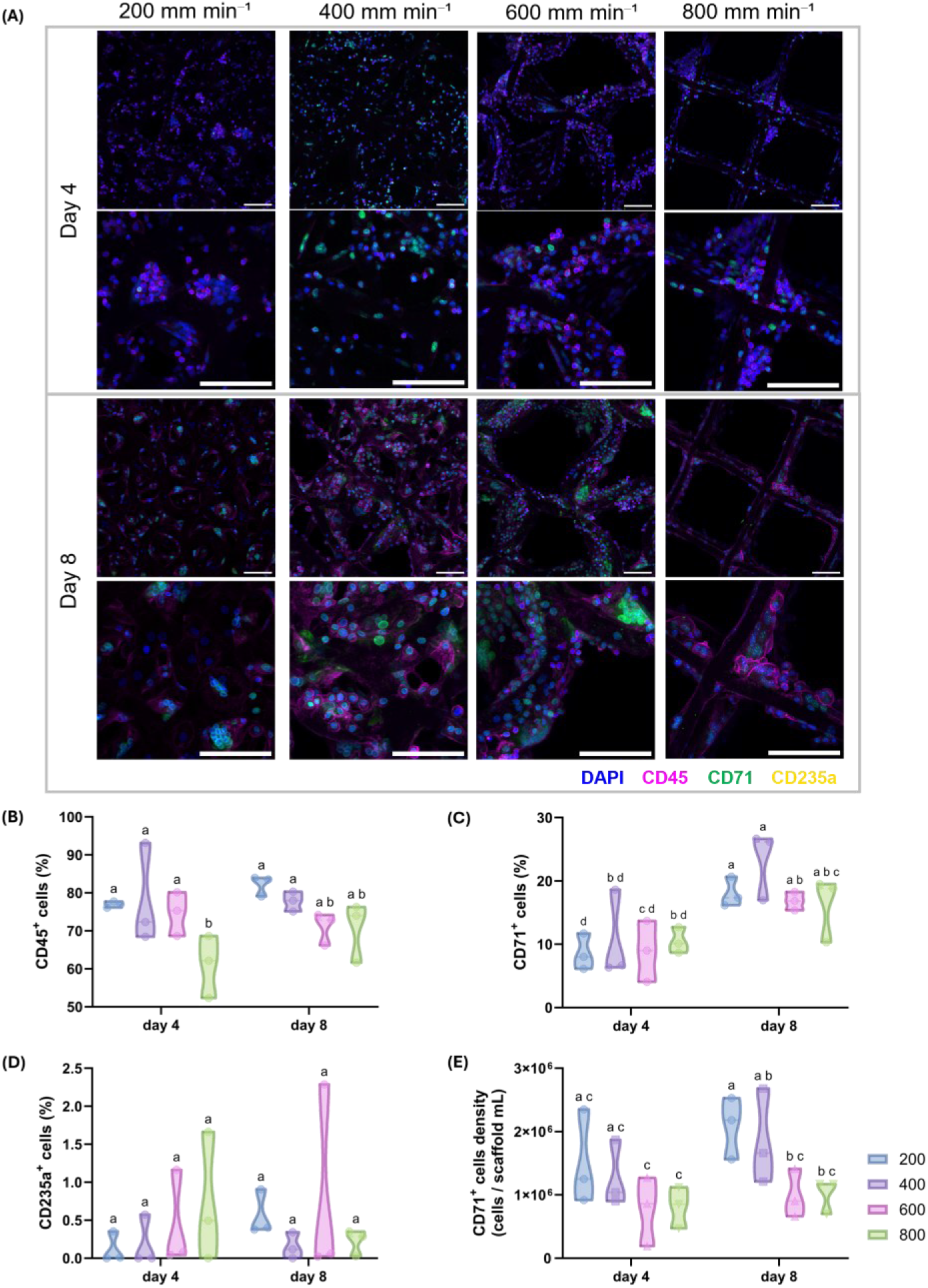
Scaffold architecture influences early erythroid differentiation of MNCs cultured on MEW scaffolds. (A) Representative maximum-intensity projections of confocal microscopy images from Donor 3 acquired after 4 and 8 days of culture on melt electrowritten (MEW) scaffolds fabricated at translational speeds of 200, 400, 600, and 800 mm min^−1^. Cells were stained with DAPI (blue), CD45 (magenta), CD71 (green), and CD235a (yellow). Scale bar = 100 µm. (B) Percentage of CD45-positive cells relative to the total number of nuclei. (C) Percentage of CD71-positive cells relative to the total number of nuclei. (D) Percentage of CD235a-positive cells relative to the total number of nuclei. (E) Density of CD71-positive cells expressed as the number of CD71-positive cells per scaffold volume (cells mm^−^3). Data were obtained from three independent donors (n = 3). Groups sharing different letters are significantly different (p < 0.05).

After colocalising each marker signal with the segmented nuclei, we quantitatively analysed the percentage and density of each cell population. Across all architectures, a progressive reduction in the proportion of CD45-positive cells was accompanied by the appearance of erythroid markers, indicating successful induction of erythroid differentiation from PB MNC cultures (figure 4 (B-D)).

Expression of transferrin receptor CD71 increased noticeably between days 4 and 8 across all scaffold architectures, indicating the emergence and expansion of committed erythroid progenitors (figure 4 (C)). CD71-positive cells were predominantly observed as dense cellular clusters occupying pore spaces between fibres and on large CD45 macrophage-like bodies rather than directly attached to scaffold surfaces. This organisation is consistent with the biological behaviour of erythroid progenitors, which are normally non-adherent and would therefore be expected to remain in suspension while becoming physically retained within the scaffold pores and interacting with the cells attached to the scaffold fibres.

In contrast to CD71, more mature CD235a-positive cells remained scarce throughout the 8-day culture period, and no clear differences were observed between scaffold architectures (figure 4 (D)), suggesting that cultures remained predominantly in the early stages of erythroid differentiation.

The 200 mm min^−1^ trabecular MEW scaffold cultures were extended to 20 days as a prototype test, using a two-part erythropoiesis medium developed by Griffiths *et al* [44]. This proof-of-concept demonstrated that a CD68 macrophage coating remained spread across MEW PCL fibres throughout 20-day erythroid culture, and the CD71 erythroblasts from days 4 and 8 differentiated into phenotypically mature CD235a red cells, which appeared to remain nucleated in erythroblast island-like macrophage clusters (figure 5). At the end of 20-day culture, significant red cell numbers detached from the scaffold and could be observed as a red pellet after centrifugation.

**Figure 5.**
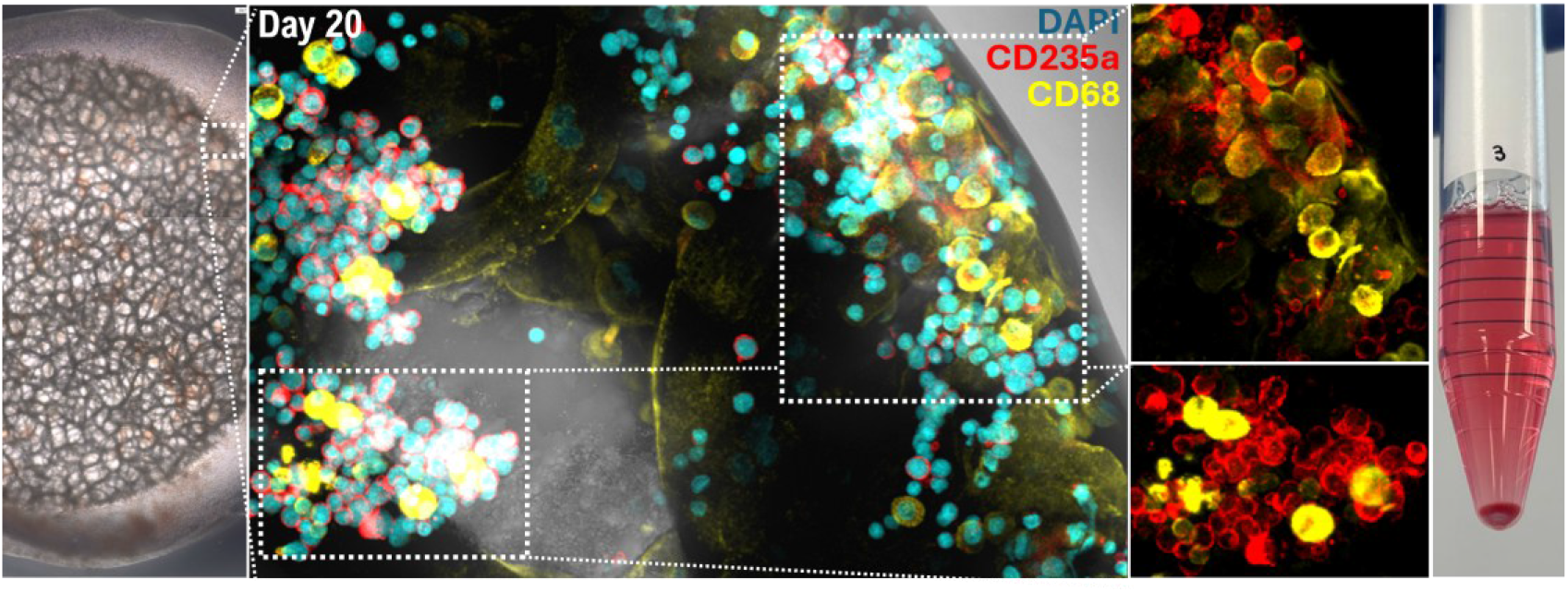
MEW scaffolds support long-term erythropoiesis to CD235a+ red cells. MEW scaffold cultures were extended to 20 days in 2-part differentiation media and contained CD68-CD235a erythroblast islands, while unattached supernatant cells formed a red pellet (right).

### 3.5. Scaffold cultures generate more supernatant cells than liquid suspension controls

All analyses presented in figures 2-4 were performed exclusively on cells attached to the scaffold at the time of fixation. However, throughout the culture period, a second cell compartment was also evident, consisting of non-adherent cells growing in suspension in the vicinity of the scaffolds (figure 6 (A)). This observation highlights an important characteristic of the proposed culture system: while scaffold fibres provide anchoring sites for MNC populations, a substantial fraction of MNCs remain in suspension. Furthermore, scaffold-attached MNCs might detach into the supernatant, and supernatant suspension cells might later attach onto the scaffold. Therefore, the MEW scaffold culture platform should be seen as a ‘supplement’ to add into conventional liquid suspension cultures, providing a macrophage feeder-layer ‘anchor’ that promotes cell-cell and cell-biomaterial interactions supportive of erythropoiesis.

**Figure 6.**
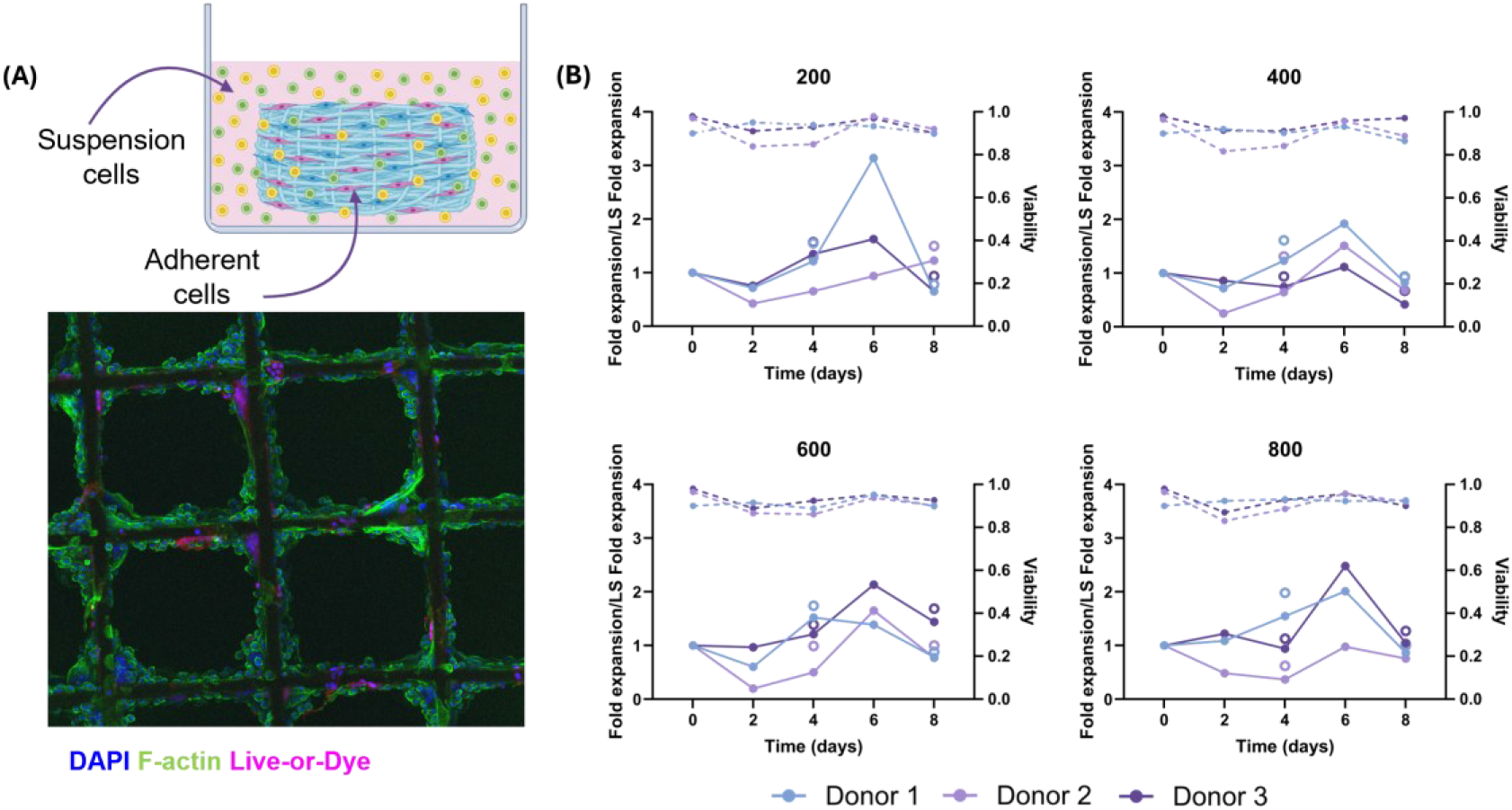
MEW scaffolds enhance supernatant cell expansion during early erythropoiesis compared to liquid suspension (LS) control. (A) MEW scaffolds were included in liquid suspension erythroid cultures, and cells grow both attached to the scaffold fibres and in the supernatant around the scaffold. Representative confocal image of cells attached to an 800 mm min^−1^ scaffold. Cells were stained with DAPI (blue), F-actin (green), and Live-or-Dye viability stain (magenta). Scale bar = 100 µm. (B). Fold expansion of liquid suspension cells growing in the proximity of MEW scaffolds printed at 200, 400, 600, and 800 mm min^−1^. Fold expansion is presented as normalised over LS fold expansion (solid lines) and cell viability (dotted lines). Empty dots represent fold expansion after including cells attached to the scaffolds. Data were obtained from three independent donors (n = 3).

Cell counting of supernatant cells revealed a common trend across all experimental groups during the early stages of culture. Both liquid suspension controls and scaffold-containing cultures experienced a substantial reduction in cell number during the first 48 h, accompanied by a decrease in viability. On average, cell numbers decreased by 92.8 ± 2.8%, while viability declined from 95.0 ± 2.7% at day 0 to 89.1 ± 2.8% at day 2. This initial loss is consistent with the selective depletion of non-adapted populations and reflects the extensive remodelling of the heterogeneous PB MNC population under erythropoietic conditions (figure S 1).

Despite this common early decline, markedly different cellular dynamics emerged thereafter. Liquid suspension controls remained largely stable after day 2 and exhibited only limited expansion throughout the remainder of the culture period. In contrast, cultures containing MEW scaffolds showed a progressive increase in cell number from day 2 onwards, peaking at day 6 before declining again by day 8 (figure 6 (B)). Although donor-to-donor variability remained substantial, all scaffold architectures displayed a tendency towards increased expansion compared with the liquid suspension controls. On average, scaffold-containing cultures achieved approximately 1.74-fold higher cell expansion than liquid suspension controls at day 6, while cultures containing the scaffold printed at 200 mm min^−1^ reached expansion values approaching 1.9-fold above the corresponding liquid suspension control.

Importantly, the enhanced expansion observed in scaffold cultures cannot be attributed solely to the contribution of scaffold-associated cells. Inclusion of the attached-cell population in the mass balance at days 4 and 8 (empty symbols in figure 6 (B)) increased the total cell yield; however, a substantial portion of the expansion advantage was already apparent in the suspension compartment alone. This finding suggests that scaffold-mediated effects are not restricted to high local densities of cells directly attached to the fibres but also extend to neighbouring cells growing in suspension. Such behaviour is consistent with the formation of local microenvironments in which scaffold-associated populations influence surrounding cells through the secretion of cytokines, growth factors, and other soluble mediators.

Interestingly, all scaffold cultures exhibited a second decline in cell number between days 6 and 8. A likely explanation is the emergence of metabolic limitations within the culture system, including depletion of essential nutrients such as glucose and/or the accumulation of metabolic by-products, particularly lactate. To investigate these possibilities, glucose consumption and lactate production were quantified by HPLC analysis of culture supernatants.

### 3.6. MEW scaffold presence increases culture metabolism and cytokine release

Metabolic analyses further demonstrate that scaffold-supported cultures differ from conventional liquid suspension systems, even if inoculated with the same number of MNCs and fed the same media volumes (figure 7).

**Figure 7.**
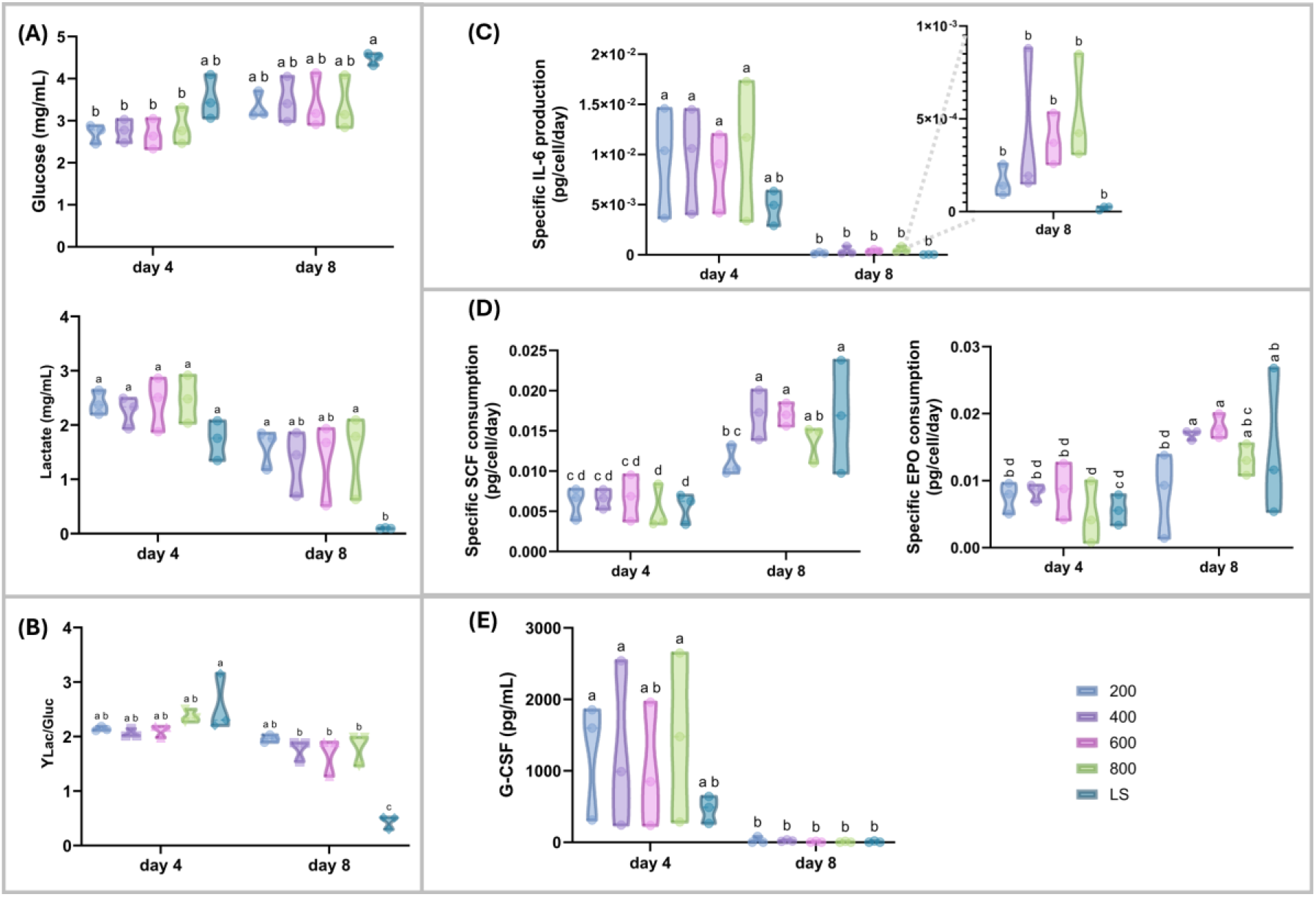
Seeding MNCs on MEW scaffolds in liquid suspension cultures affects both metabolic and cytokine dynamics during early erythropoiesis. (A) Glucose and lactate dynamics across cell culture, (B) Lactate/Glucose yield, calculated as the ratio between specific lactate production and specific glucose consumption, (C) Specific production of IL-6 as the main inflammatory marker found in culture supernatant, (D) Specific consumption of key erythropoietic growth factors: SCF and EPO, (E) G-CSF concentration measured in the culture media supernatant across culture. All specific production and consumption rates were calculated over the total cell number in the time frame analysed, not specific populations. Data were obtained from three independent donors (n = 3). Groups sharing different letters are significantly different (p < 0.05).

The limited metabolic differences observed between scaffold and liquid suspension conditions at day 4, followed by significant differences at day 8, suggest that the establishment of scaffold-associated microenvironments requires time (figure 7 (A)). Although cell attachment was already evident at an early culture time point, the results presented in figure 2-4 indicate that substantial cytoskeletal remodelling, macrophage organisation, and erythroid progenitor accumulation continue to evolve throughout the culture period. Consequently, metabolic differences between scaffold-supported cultures and liquid suspension controls may not become fully apparent until later stages of culture.

Particularly noteworthy was the marked change in the lactate-to-glucose yield observed in scaffold-supported cultures (figure 7 (B)). Because this parameter was calculated from specific glucose consumption and specific lactate production rates, it provides insight into how cells utilise available glucose rather than simply reflecting differences in cell number. Furthermore, although pH was not directly measured in the present study, increased lactate production was accompanied by colour changes of the culture medium, suggesting progressive acidification.

The substantial divergence between scaffold-supported and liquid suspension cultures indicates that scaffold inclusion fundamentally alters cellular metabolic behaviour. Given that erythroid progenitors, monocytes, and macrophages possess distinct metabolic requirements, the altered yield likely reflects differences in the cellular composition and spatial organisation established within the scaffold cultures. Although no phenotypic characterisation was performed on the liquid suspension controls, the bottom of the wells was covered with PDMS specifically included to prevent MNC attachment. Therefore, it is unlikely that significant numbers of monocytes survived and differentiated into adherent macrophages under these conditions. Consequently, the observed metabolic differences are likely to arise from the collective behaviour of the organised cellular microenvironment established around the scaffolds rather than from erythroid cells alone.

Analysing the growth factors and cytokines measured in culture supernatants, the most relevant observation was the transient inflammatory response associated with scaffold cultures. At day 4, scaffold-containing cultures produced substantially greater quantities of IL-6 than corresponding liquid suspension controls (figure 7 (C)). Across donors, IL-6 concentrations were approximately 1.5-3-fold higher in scaffold cultures compared to liquid suspension controls. A similar trend was observed for G-CSF production (figure 7 (E)). Scaffold-supported cultures produced approximately 3-5 times more G-CSF than liquid suspension controls at day 4.

Interestingly, both IL-6 and G-CSF concentrations were dramatically reduced between days 4 and 8. IL-6 concentrations decreased approximately 40-50-fold depending on donor and architecture, while G-CSF became nearly undetectable. This reduction occurred despite the persistent presence of a large CD68-positive population on the scaffold fibres. Quantitative analysis in figure 3 demonstrated that macrophage abundance remained relatively stable throughout culture and therefore cannot fully explain the observed collapse in cytokine production. Instead, these findings suggest that macrophage function changes substantially over time, transitioning from an early activated state towards a more stable scaffold-associated phenotype. The results therefore support the concept that scaffold cultures undergo an initial inflammatory adaptation phase followed by progressive establishment of a more organised microenvironment.

On the contrary, specific consumption of the erythropoietic growth factors EPO and SCF increased substantially between days 4 and 8 (figure 7 (D)). EPO consumption increased approximately two- to three-fold, while SCF consumption increased two- to four-fold depending on donor and architecture. These increases coincided with the progressive accumulation of CD71-positive erythroid progenitors observed in figure 4. Although EPO signalling is largely restricted to committed erythroid progenitors and early erythroblasts, SCF can be utilised by a broader range of haematopoietic populations, including stem cells, progenitors, myeloid populations, and erythroid precursors [51–53].

Together with the results presented in figures 2-4, these observations support a model in which MEW scaffolds function as active bioengineered microenvironments that regulate cellular organisation, cytokine dynamics, metabolism, and erythropoietic development rather than acting solely as passive structural supports.

## 4. Discussion

This study sought to engineer marrow trabeculae-mimetic MEW PCL designs supportive of cRBC biomanufacturing. Unlike advanced BM models designed to faithfully reproduce native niche physiology through the incorporation of decellularised marrow or the in vivo engineering of BM in mice [11,54], our objective was not to recreate the full complexity of the BM microenvironment. Instead, selected structural features of trabecular bone were incorporated into a synthetic and inexpensive scaffold platform without relying on animal products or biomolecules for adhesion, to determine whether they could promote macrophage retention, erythroid development, and niche-like cellular organisation.

The results obtained for scaffold microarchitecture fall within the range reported for trabecular bone that supports adult marrow haematopoiesis, including the iliac crest, vertebral bodies, proximal femur, and sternum. Porosity values reported for healthy adult trabecular bone in these anatomical sites typically range between 70 and 90%, while pore dimensions, commonly reported as trabecular separation (Tb.Sp), are generally between 300 and 1000 µm [55]. However, scaffold fibre diameters were approximately one order of magnitude smaller than the thickness of native trabeculae, which is typically reported in the range of 150-220 µm in healthy adult bone [56]. Consequently, the scaffolds provided a substantially larger surface-area-to-volume ratio than native trabecular bone while maintaining pore dimensions within the physiological range reported for marrow cavities. Importantly, all architectures maintained interconnected porosity and open networks suitable for nutrient transport and cell infiltration. Therefore, subsequent cellular responses were interpreted as consequences of altered architectural cues rather than gross failures in scaffold accessibility.

Scaffold fibres, regardless of the specific architecture, acted as effective anchoring substrates for PB MNCs without the incorporation of additional adhesion-promoting molecules. Within the MNC fraction, monocytes typically represent 10-20% [57] and rapidly undergo phenotypic changes following adhesion to surfaces, including differentiation towards macrophages [58]. The preferential localisation of CD68-positive cells along scaffold fibres was likely influenced by both scaffold-mediated effects and the biological culture conditions. Notably, IL-3 was supplemented throughout the entire culture period and has been reported to support not only erythropoiesis but also inflammation-associated monocytopoiesis [59]. Monocytes undergo pronounced morphological changes during differentiation into macrophages, including increased cell area, enhanced adhesion, and extensive actin reorganisation, processes that are not necessarily accompanied by substantial proliferation [60–63]. Despite maintaining high viability, the scaffold-associated population declined between days 4 and 8, and this could be because the cell turnover exceeds their limited capacity for self-renewal or because the available surface area in the scaffold has been occupied. The morphological changes observed were consistent with previous studies demonstrating macrophages polarisation and inflammatory activity can be controlled by MEW-generated architectures [64]

Cell densities of up to 2 × 10^7^ cells mL^−1^ were achieved within the more intricate scaffold architectures, representing a substantial increase in volumetric productivity compared with those typically reported in standard suspension cultures, in the range of 10⁵-10⁶ cells mL^−1^ [7,28,44]. Allenby et al. reported tissue-like cell densities exceeding 2 × 10⁹ cells mL^−1^ in a perfused three-dimensional bone marrow biomimicry system; however, that platform incorporated continuous perfusion, 28-day culture, and a thicker 3D scaffold better able to retain cells [8]. Another recent study reported a scalable suspension agitation culture platform for differentiating hiPSC-microcarrier aggregates into functional RBCs that were able to reach cell densities around 1.7 × 10^7^ cells mL^−1^ [65]. While the cell densities achieved here remain one-to-two orders of magnitude lower than prior perfused bioreactor scaffolds, they demonstrate how different scaffold architectures can support levels of local cellular crowding.

Within the bone marrow, HSPCs are not randomly distributed throughout the marrow cavity but are spatially organised within specialised niches associated with trabecular surfaces, the endosteum, and adjacent vascular structures [10,50,66]. These regions provide extensive cell-matrix and cell-cell interactions that contribute to the regulation of haematopoietic maintenance and lineage commitment. The stability of CD68-positive cells across all architectures and time points was combined with the progressive increase in CD71-positive progenitors, suggesting the coexistence of two distinct cellular compartments: a stable macrophage-rich adherent compartment and a developing erythroid compartment located primarily within scaffold pores.

The density of CD71-positive cells was highest within the 200 mm min^−1^ scaffolds, which were also characterised by the greatest overall cellular density. This relationship suggests that erythroid progenitor enrichment is associated with high-density cellular microenvironments rather than the availability of open pore space, similar to the highly dense erythroid colonies and erythroblast-macrophage islands found in bone marrow erythropoiesis [67].

Interestingly, scaffold architectures promoting the highest erythroid progenitor densities were not those exhibiting the highest macrophage density per unit scaffold area. Rather, compact architectures favoured the accumulation of erythroid populations, whereas highly ordered architectures promoted macrophage attachment and spreading along scaffold fibres. These findings suggest that scaffold geometry independently regulates the organisation of distinct cellular compartments within the developing microenvironment.

Despite the increasing production of CD71^+^ erythroid precursors, CD235a-positive cells remained scarce during the 8-day culture period. These findings are consistent with previous studies of ex vivo erythropoiesis showing that CD71 expression appears early during erythroid commitment, whereas substantial glycophorin-A (CD235a) expression emerges only after further maturation of erythroid precursors [18,68,69]. Allenby et al. similarly reported the predominance of CD71-positive populations during the early phases of erythropoiesis in a three-dimensional bone marrow biomimetic culture system, with more mature erythroid phenotypes becoming evident at later time points [8].

Additional phenotypic characterisation showed the constant presence of HIF-1α-positive cells in scaffold cultures. Recent evidence demonstrates that HIF-1α stabilisation in monocytes and macrophages can occur not only in response to oxygen deprivation but also following inflammatory activation and metabolic stress [70]. HIF-1α has been shown to regulate macrophage survival, migration, metabolism, and phenotypic plasticity, making it particularly relevant within developing haematopoietic microenvironments [71]. Consequently, the persistence of HIF-1α-positive cells in scaffold cultures may reflect local adaptation to the metabolically active and highly crowded scaffold-associated microenvironment, rather than solely reduced oxygen availability.

Although macrophage phenotypes were not characterised, the observed reduction in cytokine production despite stable macrophage numbers suggests that changes in activation state may have occurred during culture. Macrophages are known to exhibit considerable phenotypic plasticity in response to biophysical cues, including cell spreading, cytoskeletal tension and interactions with biomaterial surfaces, while regulating inflammatory signalling and the expression of phenotype-associated markers [61,72]. Consequently, an early pro-inflammatory activation state followed by a transition towards a more homeostatic or tissue-remodelling phenotype represents a plausible explanation for the temporal cytokine profile observed here. Furthermore, progressive extracellular matrix deposition by resident cells may have altered the local microenvironment over time, reducing the influence of the original scaffold surface and attenuating inflammatory signalling. This interpretation is consistent with the recognised ability of biomaterial-associated macrophages to respond dynamically to changes in their local niche during tissue repair and foreign body responses.

While scaffolds provide anchoring sites for MNC populations, a substantial fraction of MNCs remain in suspension. Despite donor variability, the inclusion of MEW scaffolds resulted in an increased expansion of suspension cells compared to liquid suspension controls. By day 6, cultures containing MEW scaffolds exhibited an average 1.74-fold increase in cell expansion relative to the corresponding liquid suspension controls. The decline in cell expansion observed between days 6 and 8 coincided with a rapid increase in lactate concentration and visible acidification of the culture medium. Such behaviour is consistent with the well-recognised metabolic limitations of high-density erythroid cultures, where accumulation of lactate and other metabolic by-products alters the physicochemical properties of the culture environment and ultimately restricts proliferation and differentiation. Large-scale erythroid culture platforms and advanced bone marrow biomimetic systems have identified metabolic waste accumulation and inadequate environmental control as major barriers to sustained erythropoiesis, often requiring enhanced medium exchange or continuous perfusion to maintain productivity [73,74]. Previous studies have demonstrated that erythroid expansion becomes increasingly impaired below physiological pH ranges, indicating that effective control of lactate accumulation is essential for maintaining erythropoietic performance [75]. Therefore, the decrease in expansion observed at day 8 is likely attributable to metabolic inhibition rather than loss of scaffold functionality.

Finally, analysis of cytokine production revealed that scaffold-supported cultures generate a transient cytokine burst characterised by elevated IL-6 and G-CSF production during the establishment of the scaffold-associated niche. While prolonged IL-6 signalling is generally associated with suppression of normal erythropoiesis and anaemia of chronic inflammation, transient IL-6 release can support early haematopoietic responses [10,50,76]. Importantly, the transient pattern of IL-6 production observed here differs from the persistent cytokine production typically associated with experimentally induced inflammatory responses, for example by LPS, which has shown to suppress erythropoiesis [77–79]. Although G-CSF is not considered a primary regulator of erythropoiesis in human, the elevated levels observed suggest increased activity of myeloid-derived populations during the early stages of culture establishment [80,81], and prior studies have demonstrated than in mice, G-CSF suppresses erythropoiesis [82]. This early inflammatory phase was followed by increased erythropoietic growth-factor consumption (SCF and EPO), which likely reflects not only expansion of erythroid progenitors but also the development of a complex multicellular microenvironment enriched in growth-factor-responsive cells.

Collectively, the findings of this study support a model in which MEW scaffold architecture acts as a primary regulator of microenvironment formation rather than simply providing a surface for cell attachment. Scaffold fibres promoted the preferential retention and spatial organisation of macrophage-like cells, while simultaneously supporting the accumulation of erythroid progenitors within highly populated pore-associated compartments. The coexistence of these distinct cellular populations was accompanied by transient inflammatory activation, followed by increased growth-factor consumption, altered metabolic behaviour, and enhanced cell expansion compared with conventional liquid suspension cultures. Together, these observations suggest that scaffold architecture influences erythropoiesis indirectly through the establishment of niche-like interactions between adherent and non-adherent cell populations. Although the resulting structures cannot yet be considered true erythroblastic islands, they reproduce several of their defining characteristics, including macrophage retention, localised erythroid clustering, high cellular density, and dynamic cytokine signalling. The ability to engineer these features through controlled MEW microarchitectures highlights the potential of scaffold-guided niche formation as a strategy to improve the physiological relevance and productivity of future ex vivo erythropoiesis platforms.

## 5. Conclusions

MEW fabrication enabled the generation of 2.5D scaffolds with systematically varied microarchitectures that significantly influenced cell attachment, spatial organisation, and early erythroid development. The resulting architectures promoted the establishment of highly populated cellular microenvironments, with compact scaffolds supporting the greatest overall cellular and erythroid progenitor densities, while highly ordered architectures favoured macrophage occupancy and spreading along scaffold fibres. Cultures established from minimally processed PB MNCs maintained high viability and demonstrated enhanced expansion compared with conventional liquid suspension controls, highlighting the capacity of scaffold-supported systems to influence the behaviour of both adherent and suspension cell populations.

A key finding of this study was the preferential localisation of CD68-positive macrophage-like cells on scaffold fibres, where they formed a stable adherent compartment. From day 4 to 8, CD71-positive erythroblasts grew from 10% to 25% of MNCs, which adhered in clusters against these macrophage-like cells in formations reminiscent of bone marrow erythroblastic islands. Scaffold-supported cultures also exhibited distinct metabolic and cytokine profiles, characterised by transient increases in IL-6 and G-CSF production, enhanced EPO and SCF consumption, and altered glucose and lactate utilisation. These findings indicate that scaffold architecture regulates not only cellular organisation but also the biochemical environment experienced by developing haematopoietic populations.

Altogether, the results support a model in which MEW scaffolds function as active bioengineered microenvironments rather than passive adhesion substrates. By promoting macrophage retention, local cellular crowding, and niche-like interactions, these systems partially recapitulate key features of the bone marrow microenvironment and provide a promising platform for studying early erythropoiesis. Future integration with perfusion technologies and long-term culture strategies may further enhance the physiological relevance of this approach and contribute to the development of advanced ex vivo erythropoietic manufacturing platforms

## 6 Acknowledgments

This work was supported by the Australian Research Council (DE220100757), the Ramaciotti Foundation (2022HIG08), Advance Queensland (AQIRF1312018), and the National Heart Foundation of Australia (108132-2024_FLF). This work used the Queensland node of the NCRIS-enabled Australian National Fabrication Facility (ANFF).

## Supplementary data

### 6.1. Supplemental discussion on donor viability

Increased donor-to-donor variability was observed, particularly for Donor 2, who was substantially older than the other participants (supplementary Table 1). Although no conclusions regarding ageing can be drawn from the present dataset, age-related changes in haematopoietic stem-cell function may have contributed to the reduced cellular response.

Studies of phenotypic characterisation of hematopoietic stem cells (HSPCs) have established that the number of these cells in the BM increases by two to tenfold as we age [83]. However, studies under stress and regeneration conditions, such as serial transplantation experiments, have shown that aged HSPCs have a decreased regenerative potential [84]. In conclusion, the greater number of aged HSPCs does not compensate for the loss of their functions and results in a depleted regenerative capacity.

**Table S1.** Donor variability including age, phenotype and MNC count after isolation.

| Donor | Age | Gender | Blood Type | MNC count | Other phenotypic information |
| --- | --- | --- | --- | --- | --- |
| 1 | 32 | Female | O<sup>+</sup> | 4.74 x 10<sup>8</sup> | C-K-Fyb-Jka-S- |
| 2 | 82 | Male | A<sup>+</sup> | 6.42 x 10<sup>8</sup> | C-E+c+e+K-Fya-Fyb+Jka+JKb-M-S-s+ |
| 3 | 27 | Female | O<sup>+</sup> | 4.38 x 10<sup>8</sup> | R1R2 C+E+c+e+K- |

### 6.2 Supplemental results on liquid suspension cell dynamics

The inclusion of MEW scaffolds resulted in the establishment of two distinct cellular compartments: cells attached to the scaffold fibres and cells remaining in suspension in the vicinity of the scaffold. Cell number and viability were monitored every 48 h in the suspension fraction of both scaffold-supported cultures and liquid suspension controls.

A substantial reduction in cell number was observed during the first 48 h of culture across all experimental conditions, followed by a period of relative stabilisation (figure S 1). This initial decrease is consistent with the depletion of cell populations that were not supported by the erythroid culture conditions, resulting in enrichment of cells capable of adapting to the culture environment. Despite notable donor-to-donor variability, cultures containing MEW scaffolds displayed a general tendency towards increased cell numbers between days 2 and 6 compared with the corresponding liquid suspension controls.

Following day 6, cell numbers declined in most scaffold-supported cultures and were accompanied by a reduction in cell viability. This behaviour suggests the emergence of metabolic limitations at later culture time points. Although pH was not directly measured, progressive changes in medium colour were consistently observed, indicating acidification of the culture environment. These observations are consistent with the accumulation of metabolic by-products, such as lactate, which was subsequently confirmed by HPLC analysis and may have contributed to the reduced expansion observed during the final stages of culture.

### 6.3. Supplemental methods on image analysis

Cell counting was performed at day 4 and day 8 of erythroid cultures by counting nuclei using the StarDist plugin of Fiji. It is assumed every cell contains exactly one nucleus, as PB MNCs were seeded, and erythroid enucleation would be negligible by day 8. However, differentiated macrophages which phagocytose other MNCs and become multinucleated could impair these analyses. Specifically, the TrackMate – StarDist plugin was employed in Python and Fiji (ImageJ) using the “Simple LAP Tracker.” As shown in figure S 2 (A), spot detection settings used: quality > 0.48. Quality is linked to the neural network detection confidence/probability of that segmented object. Perimeter < 51.2, based on literature references for macrophage nuclei diameter across different activation states (5 – 16 µm) [85], and assuming circularity, would represent parameters around 15 – 50 µm. Mean intensity > 76.7. The mean intensity is the average pixel value inside the segmented object. All values selected were assessed visually. StarDist nuclear segmentation was found to be 98% accurate when compared to manual scoring for a range of 8 images (from the total 120), covering the three donors, the two time points and the four architectures, as shown in figures S 2 (B-C).

**Figure S 1.**
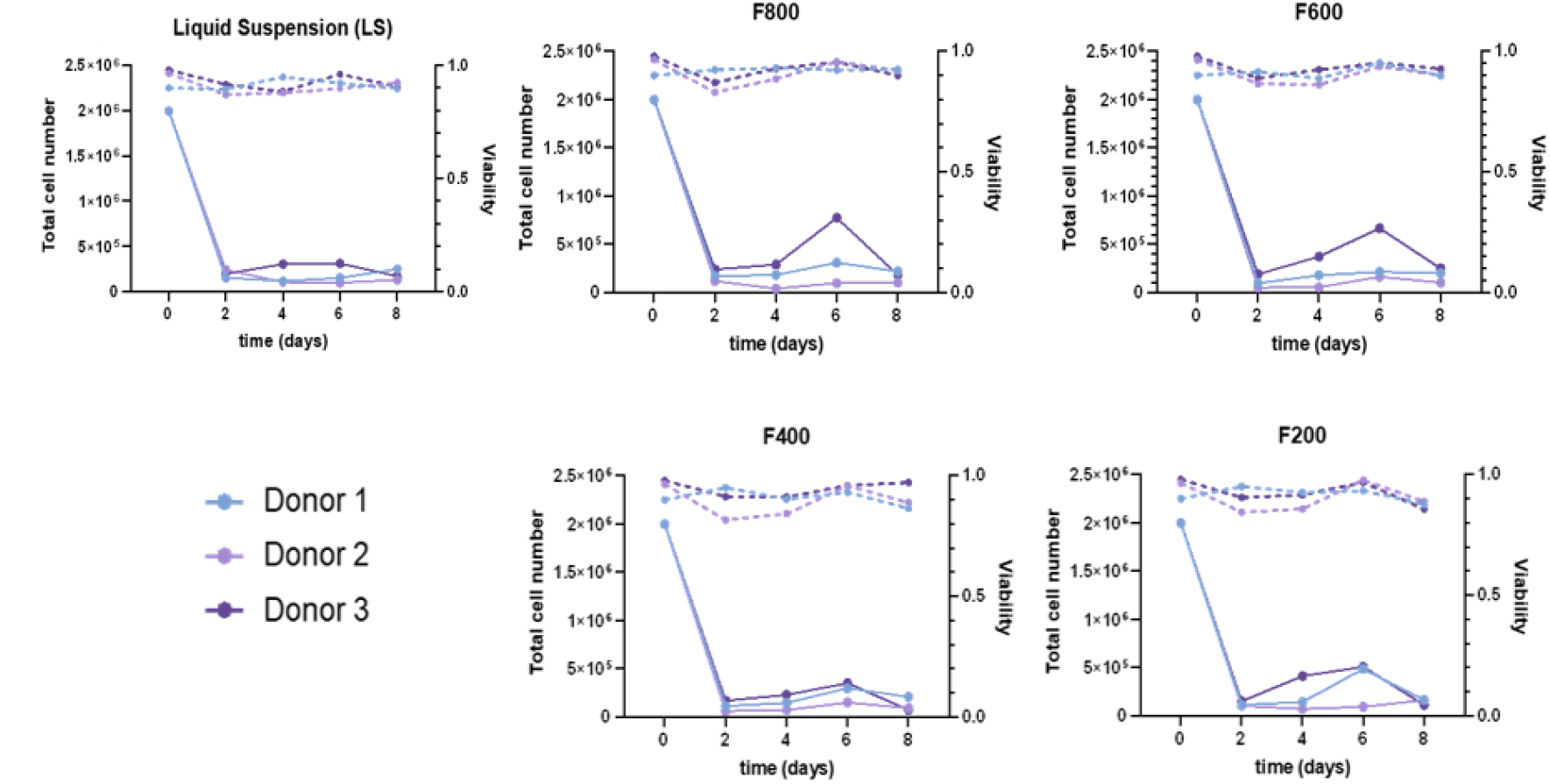
Liquid suspension cell dynamics. Total suspended cell number (solid lines) and viability (dotted lines) for liquid suspension control and cultures including scaffolds

StarDist was then similarly applied to segment phenotypic stains. The distribution of phenotypic stains CD45, CD71, and CD235a were compared to nuclear segmentations to determine each cell’s phenotype (assuming one nuclei per cell). This was done in Python by extracting DAPI segmented polygons from Fiji StarDist .xml file outputs and converting them to .csv files, defining nuclear area for each detected spot, dilating nuclear area by 5 pixels (where 1 pixel is 0.6215 µm), and then comparing stain phenotype expression within the nuclear segmentation or the 5 pixel perinuclear area. Then, a phenotype stain brightness threshold was determined by analysing isotype images similarly and calculating the mean isotype brightness for each cell’s nuclear and perinuclear area for each of the 100-2,000 nuclei detected per image and setting an isotype threshold at 99.9% for each phenotype channel. This was then similarly applied for the positive samples, where a cell was considered positive for a phenotypic stain if its mean nuclear or perinuclear phenotype brightness was above its respective 99.9% isotype threshold. figure S 3 displays representative results of this phenotype co-localisation analysis.

**Figure S 2.**
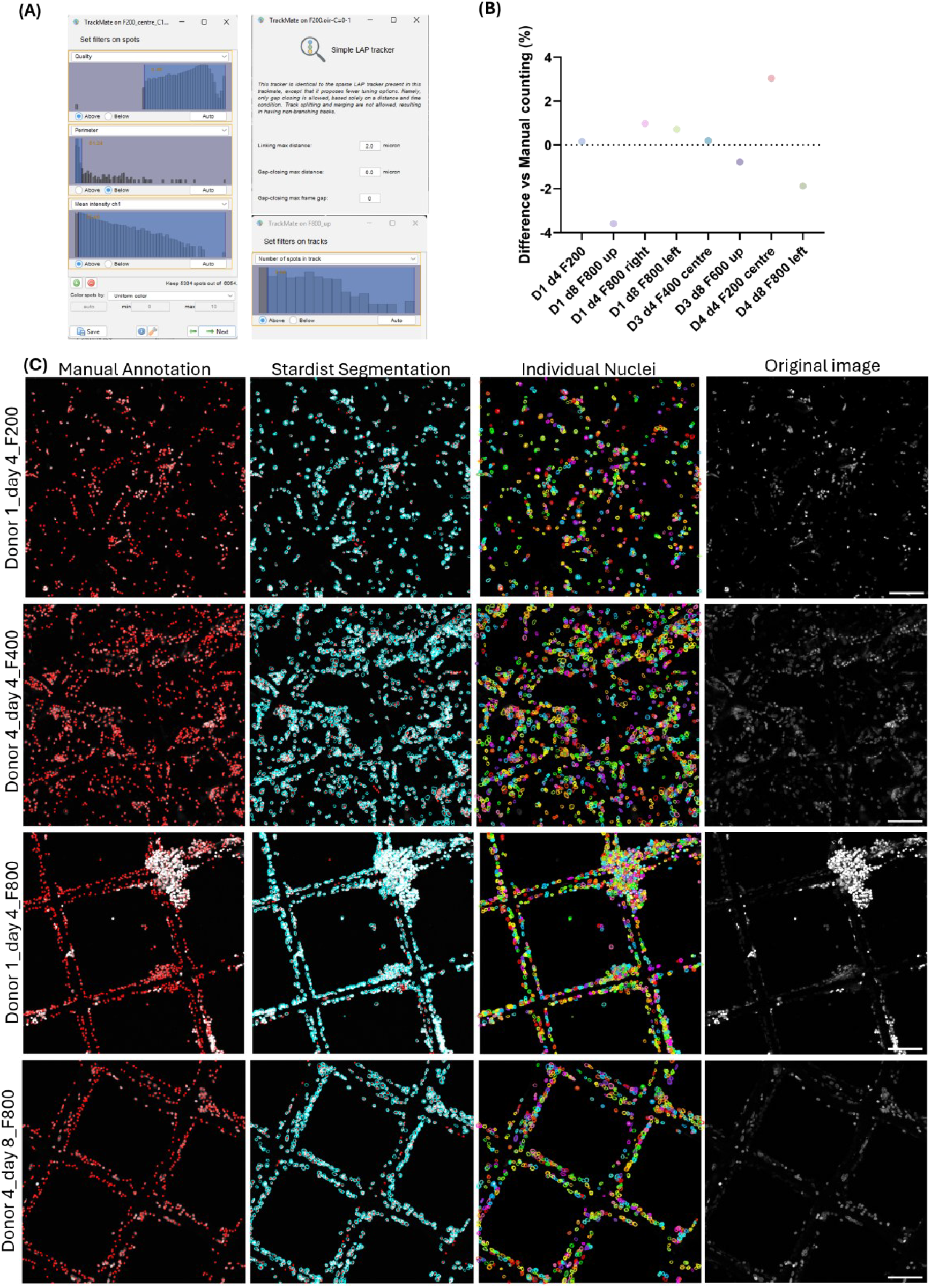
Automated cell counting by Fiji StarDist nuclear segmentation. (A) Parameters used during analyses. (B) Summary accuracy compared to manual scoring of 8 representative images. (C) Manual scoring and StarDist automated scoring of representative images. Scale bars: 100 µm.

**Figure S 3.**
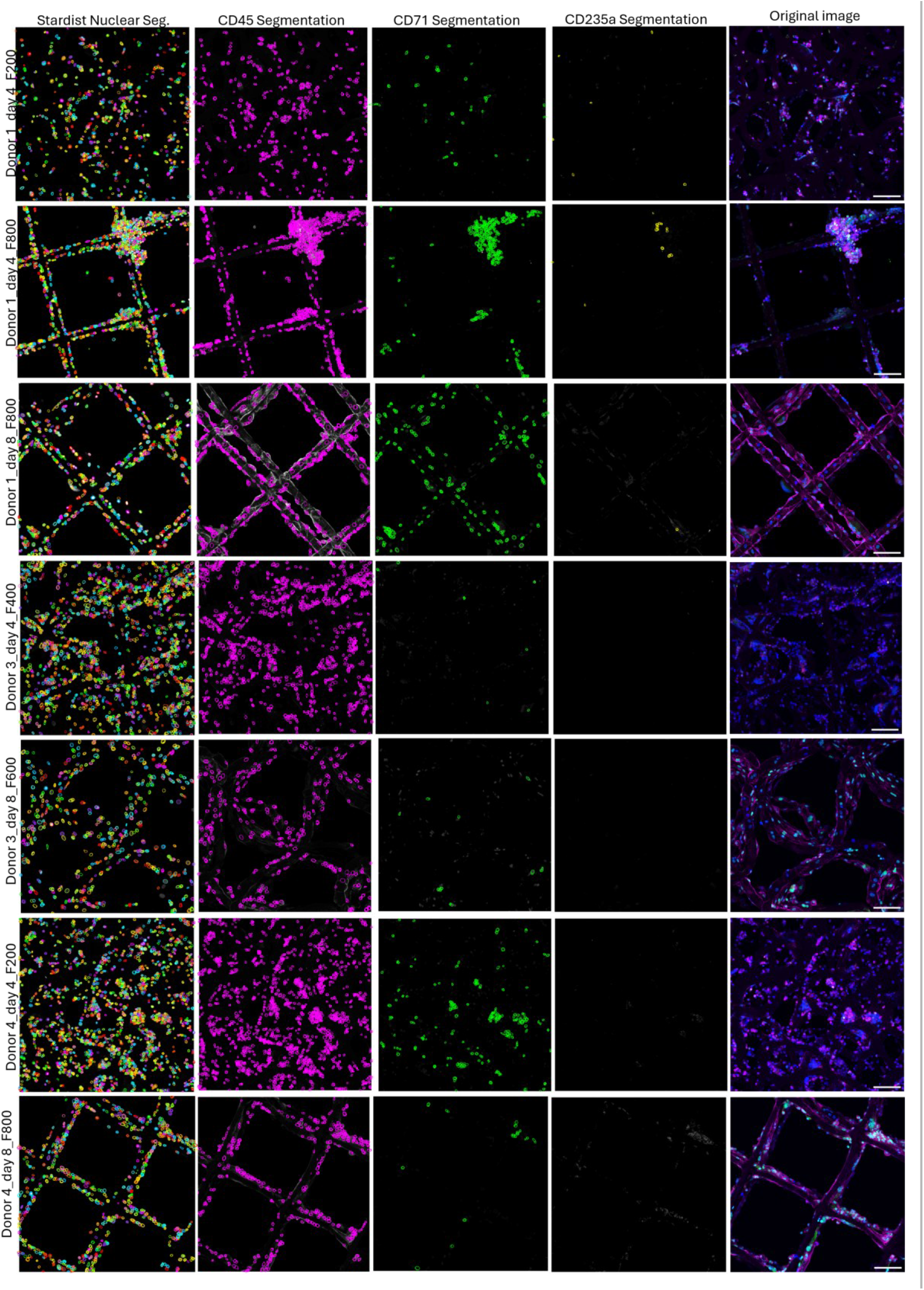
Representative nuclear and phenotypic stain segmentation by Fiji StarDist for colocalization and cell phenotyping analyses. Scale bars: 100 µm.

CD45 is a transmembrane protein tyrosine phosphatase expressed on most of the nucleated haematopoietic cells and their precursors, with the exception of mature erythrocytes and platelets [86]. As erythroid differentiation progresses, CD45 expression is progressively lost, making it a useful marker to monitor the depletion of non-erythroid and early haematopoietic populations. CD71 is the transferrin receptor responsible for iron uptake and is highly expressed in proliferating erythroid progenitors due to their elevated iron requirements for haemoglobin synthesis [87,88]. Expression typically appears during early erythroid commitment and gradually decreases as terminal maturation is approached [89,90]. CD235a (glycophorin A) is a membrane glycoprotein specifically associated with the erythroid lineage and is commonly used as a marker of more mature erythroid cells. Its expression begins during the basophilic erythroblast stage and is maintained throughout subsequent stages of erythroid differentiation [91].

### 6.4. Supplemental details on analytical methods

Glucose and lactate concentrations were determined in the culture supernatant at days 4 and 8. Metabolite identification and quantitation were done after comparison with standards and calibration curves. figure S4 showcases a representative chromatogram of one of the calibration points, one representative sample and calibration curves for peak area.

**Figure S 4.**
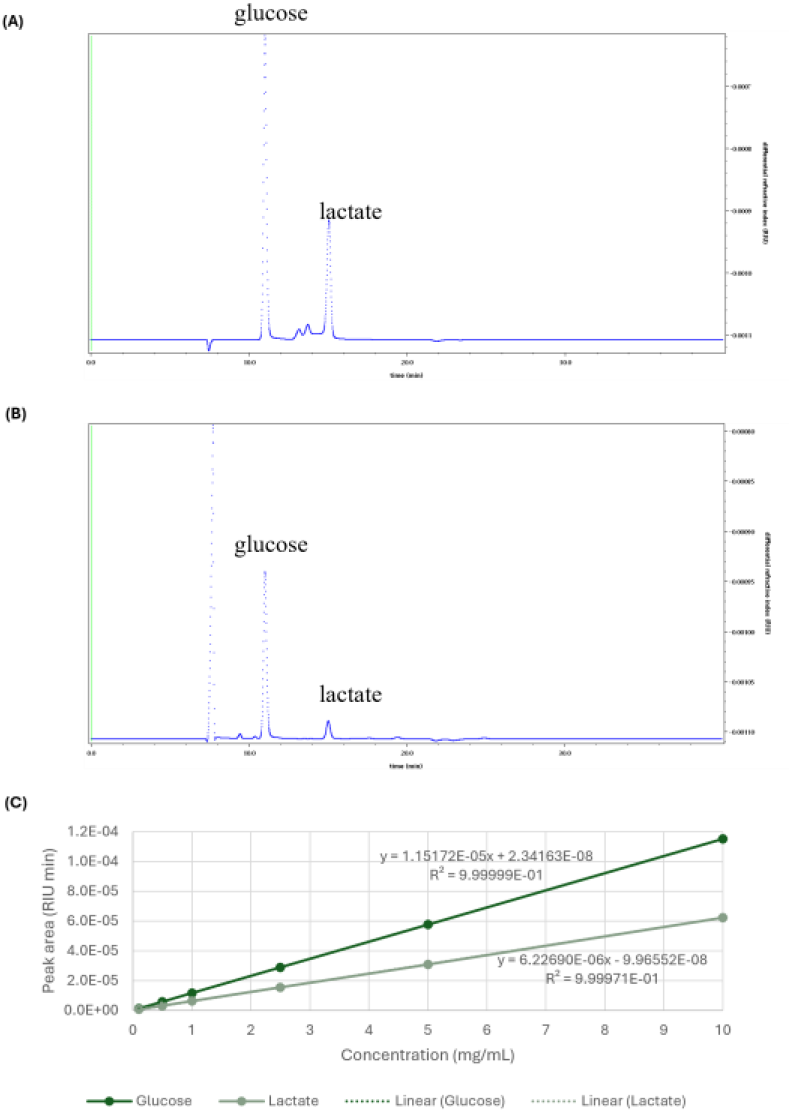
HPLC qualitative and quantitative method for determination of glucose and lactate in culture medium supernatants. (A) Chromatogram for 10 mg mL^−1^ standards (B) Representative chromatogram of a sample (Donor 2 day 8 F200). (C) Calibration curves for peak area including standards with 0.1, 0.5, 1.0, 2.5, 5, and 10 mg mL^−1^.

Mix and match LEGENDplex™ kits were used for the quantitation of key growth factors (SCF, EPO, G-CSF, and PDGF-AA) and inflammation markers (TNF-α, IFN-γ, and IL-6). Figures S 5-6 include the calibration curves and supernatant concentrations measured for every analyte.

**Figure S 5.**
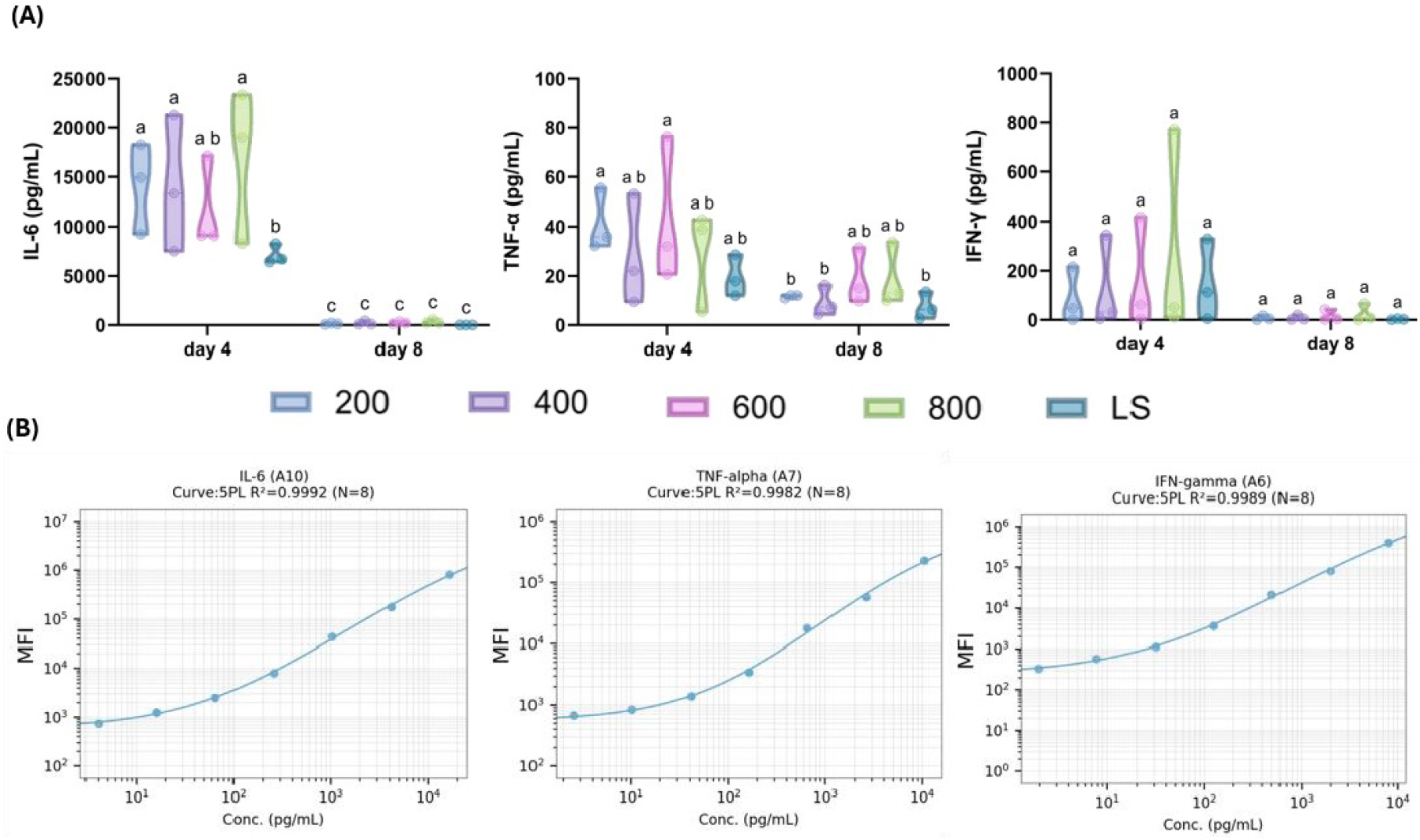
LEGENDplex analysis for inflammatory markers. (A) Cytokine concentrations measured in cell culture supenatant for IL-6, TNF-α, and IFN-*γ*. (B) Calibration curves for LEGENDplex analysis

**Figure S 6.**
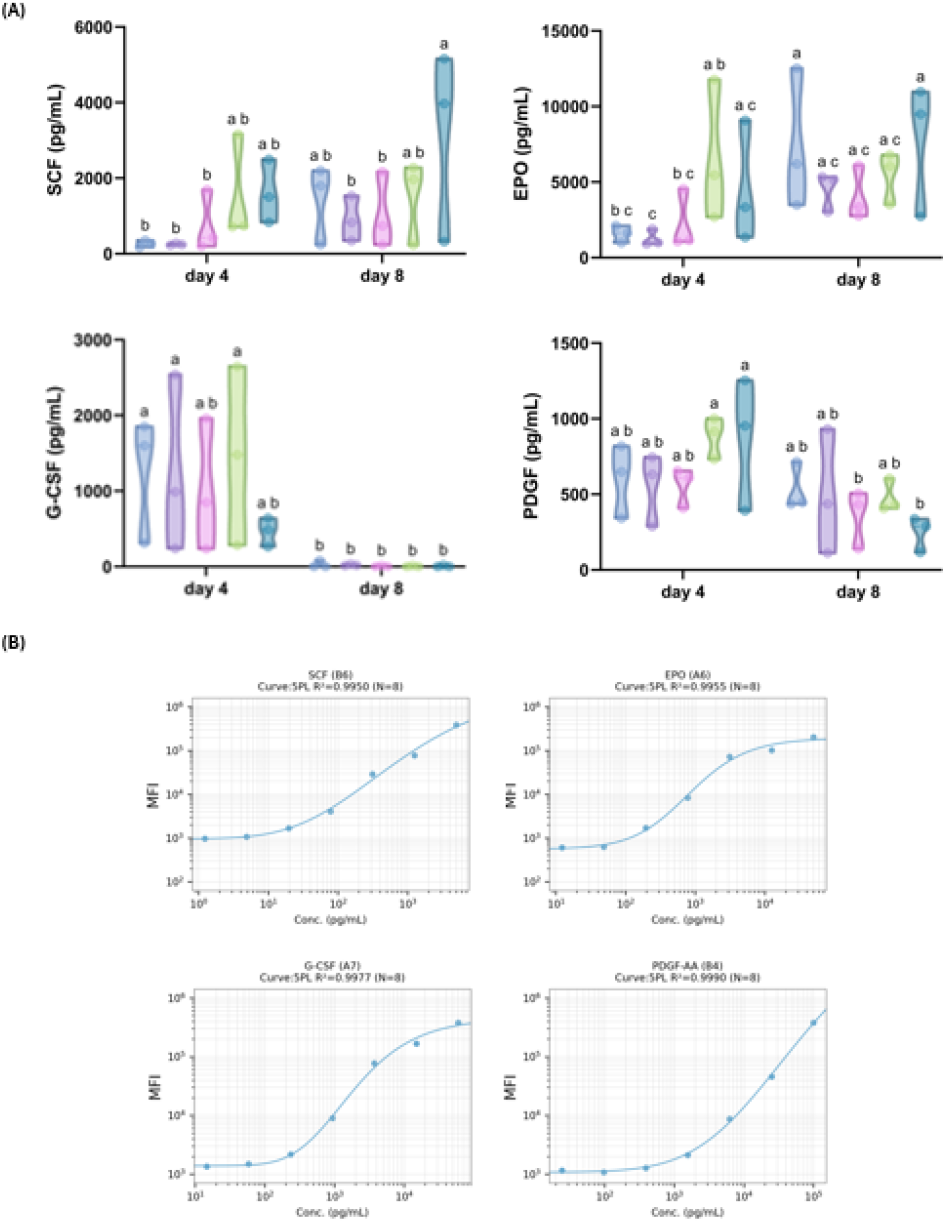
LEGENDplex analysis for growth factors. (A) Cytokine concentrations measured in cell culture supernatant for SCF, EPO, G-CSF, PDGF. (B) Calibration curves for LEGENDplex analysis

